# Genes vary greatly in their propensity for collateral fitness effects of mutations

**DOI:** 10.1101/2022.10.24.513589

**Authors:** Jacob D. Mehlhoff, Marc Ostermeier

## Abstract

Mutations can have deleterious fitness effects when they decrease protein specific activity or decrease active protein abundance. Mutations will also be deleterious when they cause misfolding or misinteractions that are toxic to the cell (i.e., independent of whether the mutations affect specific activity and abundance). The extent to which protein evolution is shaped by these and other collateral fitness effects is unclear in part because little is known of their frequency and magnitude. Using deep mutational scanning (DMS), we previously found at least 42% of missense mutations in the *TEM-1* β-lactamase antibiotic resistance gene cause deleterious collateral fitness effects. Here, we used DMS to comprehensively determine the collateral fitness effects of missense mutations in three genes encoding the antibiotic resistance proteins New Delhi metallo-β-lactamase (NDM-1), chloramphenicol acetyltransferase I (CAT-I), and 2”-aminoglycoside nucleotidyltransferase (AadB). *AadB* (20%), *CAT-I* (0.9%), and *NDM-1 (*0.2%) were less susceptible to deleterious collateral fitness effects than *TEM-1* (42%) indicating that genes have different propensities for these effects. As was observed with *TEM-1*, all the studied deleterious *aadB* mutants increased aggregation. However, aggregation did not correlate with collateral fitness effects for many of the deleterious mutants of *CAT-I* and *NDM-1*. Select deleterious mutants caused unexpected phenotypes to emerge. The introduction of internal start codons in *CAT-1* caused loss of the episome and a mutation in *aadB* made its cognate antibiotic essential for growth. Our study illustrates how the complexity of the cell provides a rich environment for collateral fitness effects and new phenotypes to emerge.

## Main Text

The evolution of proteins relies on an interplay between mutations and selective pressure. Mutations with beneficial or neutral fitness effects persist, contributing to genetic diversity, while mutations with deleterious fitness effects are filtered out of the population with time. Understanding the mechanisms underlying the fitness effects of mutations is important for the study of evolution. Building on the observations that protein misfolding and misinteractions can cause deleterious fitness effects (Parsell and Sauer 1989; Krylov et al. 2003; Lemos et al. 2005; Geiler-Samerotte et al. 2011; Olzscha et al. 2011; Levy et al. 2012; Yang et al. 2012; Navarro et al. 2014; Bratulic et al. 2015), we have proposed that it is useful to broadly categorize the fitness effect of mutations into two types: primary and collateral (Mehlhoff et al. 2020). Primary fitness effects are those caused by changes in the ability of the gene to perform its physiological function and arise from changes in protein activity and protein abundance. Collateral fitness effects are those that do not derive from changes in the gene’s ability to perform its physiological function. Misinteractions, whether from proteins in a misfolded or natively folded state, are a likely source of deleterious collateral fitness effects. Primary and collateral fitness effects are not mutually exclusive. Mutational effects may be only primary, only collateral, or a combination of both. Knowledge of the frequency and mechanisms by which collateral fitness effects arise is essential to understanding evolutionary pathways and constraints. For example, protein misfolding and misinteractions have been proposed as reasons for the E-R anticorrelation (Zhang and Yang 2015) – the observation that protein evolution rates are best predicted by an anti-correlation with protein expression levels – but their roles continue to be debated (Biesiadecka et al. 2020; Usmanova et al. 2021; Wu et al. 2022). Under the protein misfolding and misinteraction hypotheses, highly expressed proteins evolve slower because high protein levels make protein misfolding and misinteractions more likely and more significant (i.e. both protein misfolding and misinteractions increase with concentration), and misfolding and misinteractions tend to cause negative fitness effects. In addition, while collateral fitness effects are possible whenever the gene is expressed, primary fitness effects only manifest when the gene is expressed, is under selective pressure for its physiological function, and is not in a regime where fitness is buffered against the mutational effects on protein function and abundance (Mehlhoff et al. 2020).

Collateral fitness effects of mutations can be observed by growing cells in environments in which the physiological function(s) of the gene are irrelevant to organismal fitness. Studying the fitness effect of mutations in antibiotic resistance genes in environments lacking their relevant antibiotic is a convenient model system because any fitness change can be presumed to be a collateral fitness effect. We previously measured the collateral fitness effects for all single-codon missense and nonsense mutations in the antibiotic resistance gene *TEM-1* beta-lactamase (Mehlhoff et al. 2020). TEM-1 is a periplasmic protein that is exported with its N-terminal signal sequence via the Sec export pathway. Upon crossing the inner membrane during export, the signal sequence of TEM-1 is cleaved by signal peptidase I. The mature TEM-1 protein then folds via intermediates in which the α domain of the protein (residues 69 to 212) is collapsed followed by the assembly of the two lobes of the α/β domain. One surprising finding from our study was that deleterious collateral fitness effects were very common. At least 42% of missense mutations caused a 1% or greater fitness reduction. Mutations causing deleterious collateral fitness effects included signal sequence mutations that caused improper signal sequence cleavage and aggregation, mutations involving cysteine that caused incorrect intermolecular disulfides and aggregation, nonsense mutations that caused truncated proteins to aggregate, and mutations in the α domain that caused aggregation. Mutations causing deleterious collateral fitness effects caused activation of select outer-envelope stress pathways, but not all mutations activated the same pathways, suggesting different mechanisms for the deleterious effects.

We wondered if the high frequency of deleterious collateral fitness effects in *TEM-1* was typical of most genes. We also wondered if aggregation would be associated with the deleterious collateral fitness effects in other genes. Here, we report the frequency and magnitude of collateral fitness effects of mutations in three different bacterial antibiotic resistance genes native to *E. coli*. We explore the mechanistic determinants of these effects including the role of aggregation and describe unexpected phenotypes caused by some mutations.

## Results

### Deleterious collateral fitness effects of mutations occur less frequently in NDM-1, CAT-I, and AadB relative to TEM-1

We expanded our study of collateral fitness effects of mutations by choosing proteins that differed from TEM-1 by quaternary structure, subcellular location, and post-translational processing. We initially identified New Delhi metallo-β-lactamase (*NDM-1*), chloramphenicol acetyltransferase I (*CAT-I*), 2”-aminoglycoside nucleotidyltransferase (*aadB*), and aminoglycoside 6’-N-acetyltransferase (*aac(6’)-Im*) as candidate antibiotic resistance genes for studying collateral fitness effects. All four genes have no known physiological functions outside of providing antibiotic resistance, making the genes superfluous during growth in the absence of antibiotic.

NDM-1 is a class B metallo-β-lactamase which has been natively found in *E. coli* as a 270-residue protein and has undergone rapid, worldwide dissemination since being first identified in 2008 (Yong et al. 2009). NDM-1 evolved independently from TEM-1 and relies on two zinc ions in the active site to coordinate a nucleophilic attack of β-lactam antibiotics including penicillins, cephalosporins, and carbapenems (Zalucki et al. 2020). NDM-1 offered a periplasmic protein to compare with TEM-1. NDM-1 is reported to exist as a monomer (Guo et al. 2011) or a mixture of monomers and dimers in solution (King and Strynadka 2011). NDM-1 contains two cysteines (C26 and C208) but unlike TEM-1 does not form a native disulfide bond. Several studies have shown that it anchors to the outer membrane following export to the periplasm via the Sec export pathway, cleavage of its signal peptide, and lipidation at C26 (González et al. 2016; Prunotto et al. 2020). However, NDM-1’s categorization as a lipoprotein has been challenged (Zalucki et al. 2020).

The other three proteins were cytoplasmic proteins. Chloramphenicol acetyltransferases (CAT) provide high-level chloramphenicol resistance and are commonly divided into three types: CAT-I, CAT-II, and CAT-III (Biswas et al. 2012). Of the three variants, CAT-I is the most prevalent and native to *E. coli* (Awad et al. 2016). AadB and AAC(6’) are monomeric enzymes that covalently modify aminoglycoside antibiotics at different sites (Wright and Serpersu 2005; Cox et al. 2015). CAT-I, AadB, and AAC(6’) lack the potential for two types of mechanisms associated with deleterious effects in TEM-1 because cytoplasmic proteins lack export signal sequences and cannot form disulfides due to the cytoplasm’s reducing environment. However, the cytoplasm offers a larger number of proteins with which these proteins might misinteract. In addition, CAT-I’s homotrimer state adds oligomerization as an additional process through which collateral fitness effects might arise.

We replaced the *TEM-1* gene in pSKunk-1 with *NDM-1*, *CAT-I*, *aadB*, and *aac(6’)-Im*, placing the genes under control of the isopropyl β-D-1-thiogalactopyranoside (IPTG) inducible *tac* promoter. All four genes provided the expected antibiotic resistance in cultures containing IPTG (**supplementary fig. S1A**, Supplementary Material online). With the exception of AAC(6’), expression of the antibiotic resistance proteins caused little, if any, fitness effects (**supplementary fig. S1B**, Supplementary Material online). We eliminated AAC(6’) as a candidate for studying collateral fitness effects because of its large fitness cost of expression. Expression levels of wild-type NDM-1, CAT-I, and AadB were moderate and did not cause significant changes in the *E. coli* transcriptome (**supplementary fig. S2**, Supplementary Material online).

We constructed libraries designed to contain all possible single-codon substitutions in *NDM-1*, *CAT-I*, and *aadB* and transformed these libraries into NEB 5-alpha LacI^q^ *E. coli* cells. We used a growth competition experiment in which uninduced, exponentially growing cultures were diluted to a starting optical density (OD-600 nm) of 0.02 in lysogeny broth (LB) media. We added IPTG to induce expression and allowed the library cultures to grow for approximately 10 generations at 37°C in the presence of the plasmid maintenance antibiotic spectinomycin but in the absence of the gene’s cognate antibiotic. We thereby assumed any fitness effects of mutations measured during growth competition in the absence of the antibiotic are collateral fitness effects. We used deep sequencing to determine the frequencies of wild-type and mutant alleles at the start and end of the 10 generations of growth. We also combined the counts of synonymous alleles to determine the frequencies of missense and nonsense mutations (i.e., on the amino acid level). We calculated the fitness (*w*) from these frequencies. In doing so, our measure of fitness corresponds to the mean growth rate of cells containing a mutant allele relative to the mean growth rate of cells containing the wild-type allele. From the collateral fitness effects, we calculated a selection coefficient, *s* = *w* − 1.0, for each mutation.

In two biological replicates, we measured the collateral fitness effects for 98.6% (5322/5400), 99.9% (4375/4380), and 94.9% (3359/3540) of the possible nonsynonymous mutations in *NDM-1*, *CAT-I*, and *aadB*, respectively. The apparent beneficial fitness effects of nearly all mutations (including nonsense mutations) at codons 26-28 and 132-134 of *NDM-1* and codons 155-156 of *aadB* were deemed spurious for several reasons (**supplementary text and figs. S3 and S4**, Supplementary Material online) including the fact that monoculture growth experiments for select mutations at these positions confirmed they lacked any significant fitness effect (**supplementary fig. S5**, Supplementary Material online). Deleterious collateral fitness effects occurred at a much lower frequency in *NDM-1* (0.24%), *CAT-I* (0.87%), and *aadB* (3.76%) than in *TEM-1* (21.6%) using the criteria of *P*<0.01 (*Z*-test) in both replicates (**fig. 1A-D**). This method identifies the specific mutations with statistically significant deleterious effects in both replica experiments.

**Fig. 1.**
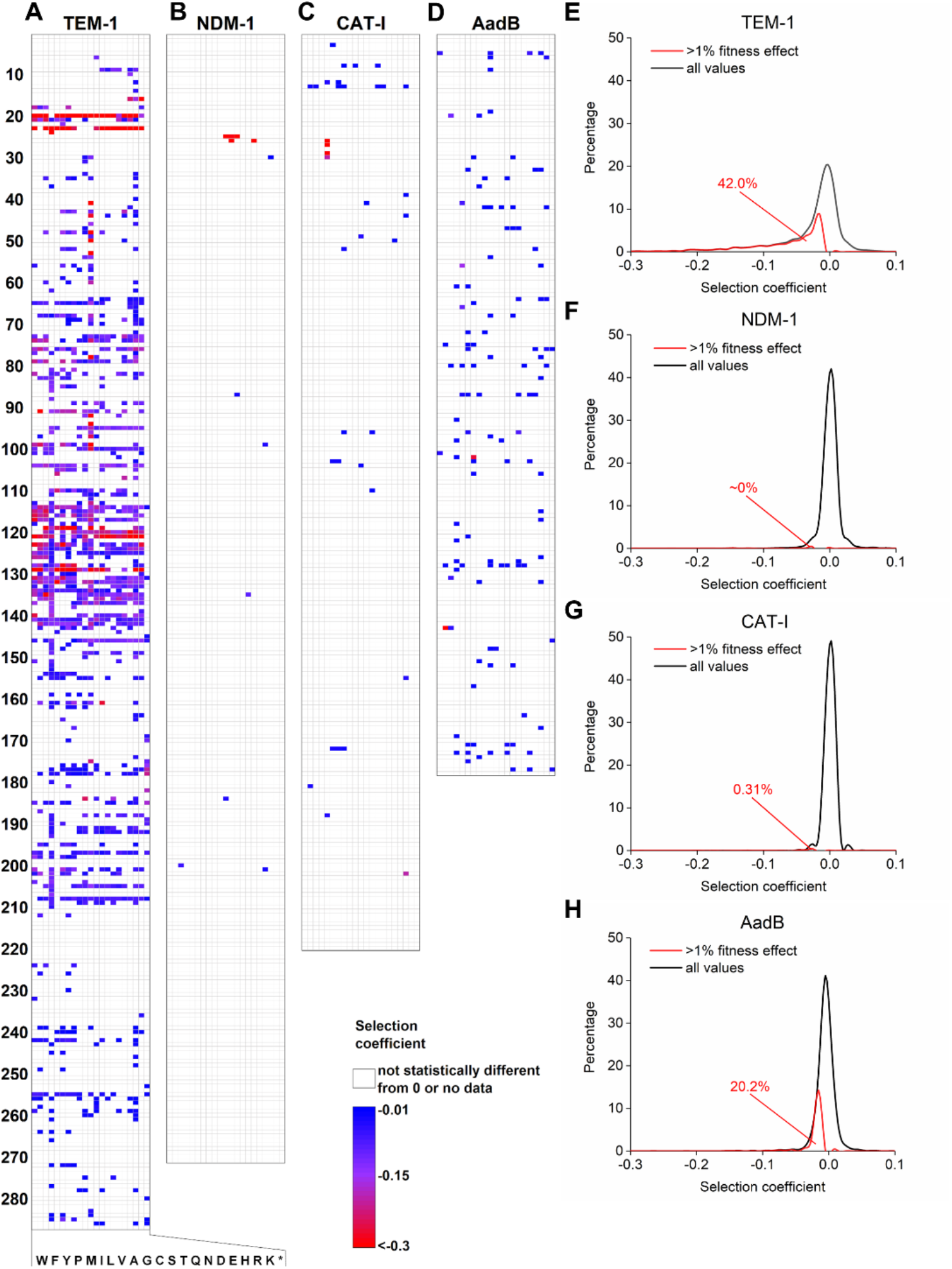
Deleterious collateral fitness effects of nonsynonymous mutations in TEM-1, NDM-1, CAT-I, and AadB. Heat map of weighted mean selection coefficients for mutations that caused significant deleterious collateral fitness effects (*P*<0.01) in both replica experiments in (**A**) TEM-1, (**B**) NDM-1, (**C**) CAT-I, and (**D**) AadB. TEM-1 data is from (Mehlhoff et al. 2020). Detailed heat maps showing all fitness values for NDM-1, CAT-I, and AadB are available as **supplementary figs. S6, S7, S8**, Supplementary Material online. DFE showing collateral fitness effects for missense substitutions in (**E**) TEM-1, (**F**) NDM-1, (**G**) CAT-I, and (**H**) AadB. The red curve and percentage indicate the estimated portion of missense mutations that cause a greater than 1% decrease in fitness by analyzing the symmetry of the distribution about the value of zero.

We also used a second method to estimate the frequency of deleterious mutations. We examined the asymmetry about a selection coefficient of 0.0 in the distribution of the weighted mean selection coefficients calculated from the two replicas. We estimated the percentage of missense mutations which caused more than a 1% decrease in fitness by assuming that selection coefficients for neutral mutations will be symmetrically distributed around a selection coefficient of 0.0 and that no mutations cause beneficial collateral fitness effects. In other words, we took the distribution of positive selection coefficients and subtracted its mirror image from the negative selection coefficient distribution. The fraction of the remaining distribution that had a value <-0.01 we deemed deleterious. This analysis confirmed that deleterious collateral fitness effects were rare in *NDM-1* and *CAT-I*, but a significant fraction of mutations in *aadB* (20.2%) have deleterious effects of >1% by this analysis (**fig. 1E-H**). The frequency of deleterious mutations in *aadB* is much higher by this method because there is a large fraction of mutations in *aadB* with small effects – too small to be deemed deleterious individually by the first method unless they happened to be highly represented in the population before the growth period.

### Confirmation of fitness effects by monoculture growth experiments

For each protein, we constructed a set of representative mutations to confirm their fitness effects using monoculture growth experiments. We monitored the growth rate for monocultures of cells expressing these mutants relative to cells expressing the wild-type genes over 10 generations of exponential growth. From the starting and final optical density (OD) of the monocultures, we calculated a mean growth rate to determine the fitness effects of these mutations (**supplementary fig. S5**, Supplementary Material online).

For *NDM-1*, only 0.24% (12/5057) of missense mutations resulted in measurable deleterious collateral fitness effects in both replica DMS experiments (*P*<0.01, *Z*-test) (**fig. 2A**). Collateral fitness effects were not observed at residues H120, H122, D124, H189, C208, and H250 responsible for coordinating the zinc ions and forming the active site. We created the five missense mutants of *NDM-1* that caused over a 25% reduction in fitness as measured by DMS (**fig. 2A**). We also constructed neutral mutants C26A and C26L (to compare with the deleterious mutations C26D and C26S) and P28A, the mutation in the naturally occurring variant *NDM-2*. The deleterious and neutral effects of all eight mutations were confirmed via monoculture growth (**fig. 2C**), but irregularities in the growth behavior of cells expressing the C26S and C26D mutants (**supplementary fig. S10A**, Supplementary Material online) led us to discover that insertion of an IS4-like element ISVsa5 family transposase within *NDM-1* would occur during the 10 generations of induced monoculture growth, complicating fitness measurements for cells expressing these two mutants. This transposase preferentially targets 5’-GCTNAGC-3’ sites (Halling and Kleckner 1982). Such a site overlaps codons 22-24 in *NDM-1*, immediately preceding where the transposase insertion often took place. The transposition evidently alleviated the deleterious collateral fitness effects of these mutations by preventing NDM-1 expression. Due to C26S and C26D’s large deleterious effect on growth, cells containing the transposition became a significant fraction of the population by generation 10 (**supplementary fig. S10B**, Supplementary Material online). We made synonymous mutations at residues 23 and 24 to change the sequence to 5’-GTTATCA-3’ to get a truer estimate of the collateral fitness effect of the C26S and C26D mutations. These synonymous mutations greatly reduced and delayed the appearance of cells with the transposition in *NDM-1* (**supplementary fig. S10C**, Supplementary Material online). Subsequent experiments examining the effects of C26S and C26D used *NDM-1* genes with these synonymous substitutions and confirmed their large deleterious effect (*s*~ −0.7) (**fig. 2C** and **supplementary fig. S10D, S10E**, Supplementary Material online).

**Fig. 2.**
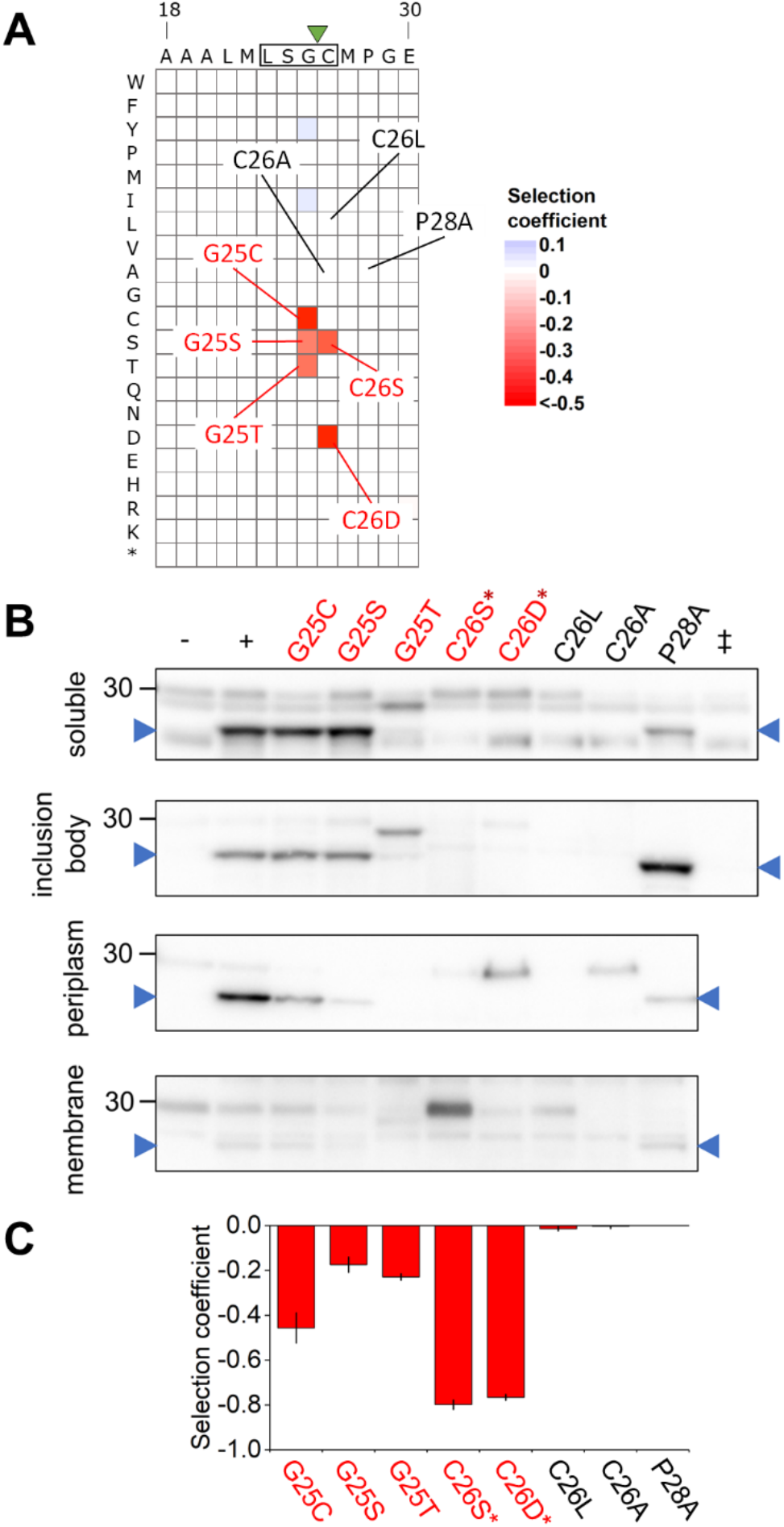
NDM-1 mutations’ effect on fitness and protein fate. (**A**) Subsection of the NDM-1 collateral fitness landscape showing weighted mean selection coefficients (by DMS) for selected mutations (*P*<0.01 in both replicates). The LSGC lipobox is indicated with a rectangle. The expected NDM-1 signal sequence cleavage by signal peptidase II site is shown by a green triangle. (**B**) Western blots of the total soluble, inclusion body, periplasmic, and membrane fractions from cells expressing unmutated NDM-1 (–, without IPTG; +, with IPTG) or the indicated NDM-1 mutants (deleterious mutations in red and neutral mutations in black). The C26S* and C26D* mutants include two synonymous mutations to prevent transposase insertion (SI Appendix, Supplementary Text). Samples from cells expressing CAT-I (‡) were included to identify the bands of cross-reacting *E. coli* proteins. The blue arrowhead indicates the expected position of mature NDM-1. Full western blots including the second biological replicate are provided as **supplementary fig. S9**, Supplementary Material online. (**C**) Selection coefficients were determined from monoculture growth experiments by measuring the mean growth rate over six hours of growth post-induction. Error bars represent 99% confidence intervals (*n*=3).

A total of 0.87% (36/4156) of missense mutations in CAT-I caused deleterious collateral fitness effects in both replica experiments (*P*<0.01, *Z*-test) (**fig. 3A**). Collateral fitness effects were not observed at key active site residues, including S146 and H193. We constructed a set of 12 missense mutations in CAT-I. Eight were chosen as a broad representation of mutations with significant deleterious effects in both replicas (including two of the stretch of four large-effect deleterious mutations to Met at positions 27-30), three were chosen because they had potential beneficial or deleterious effects based on one of the two replica experiments, and G202K was chosen as control neutral mutation for the highly deleterious G202R. Of the eight that were deleterious in both DMS data sets, seven were deleterious in the monoculture growth (**fig. 3C** and **supplementary fig. S5B**, Supplementary Material online). The three mutations with potential fitness effects based on just one replica did not show fitness effects in monoculture growth (**supplementary fig. S5B**, Supplementary Material online). G202K was neutral as expected (**fig. 3C)**.

**Fig. 3.**
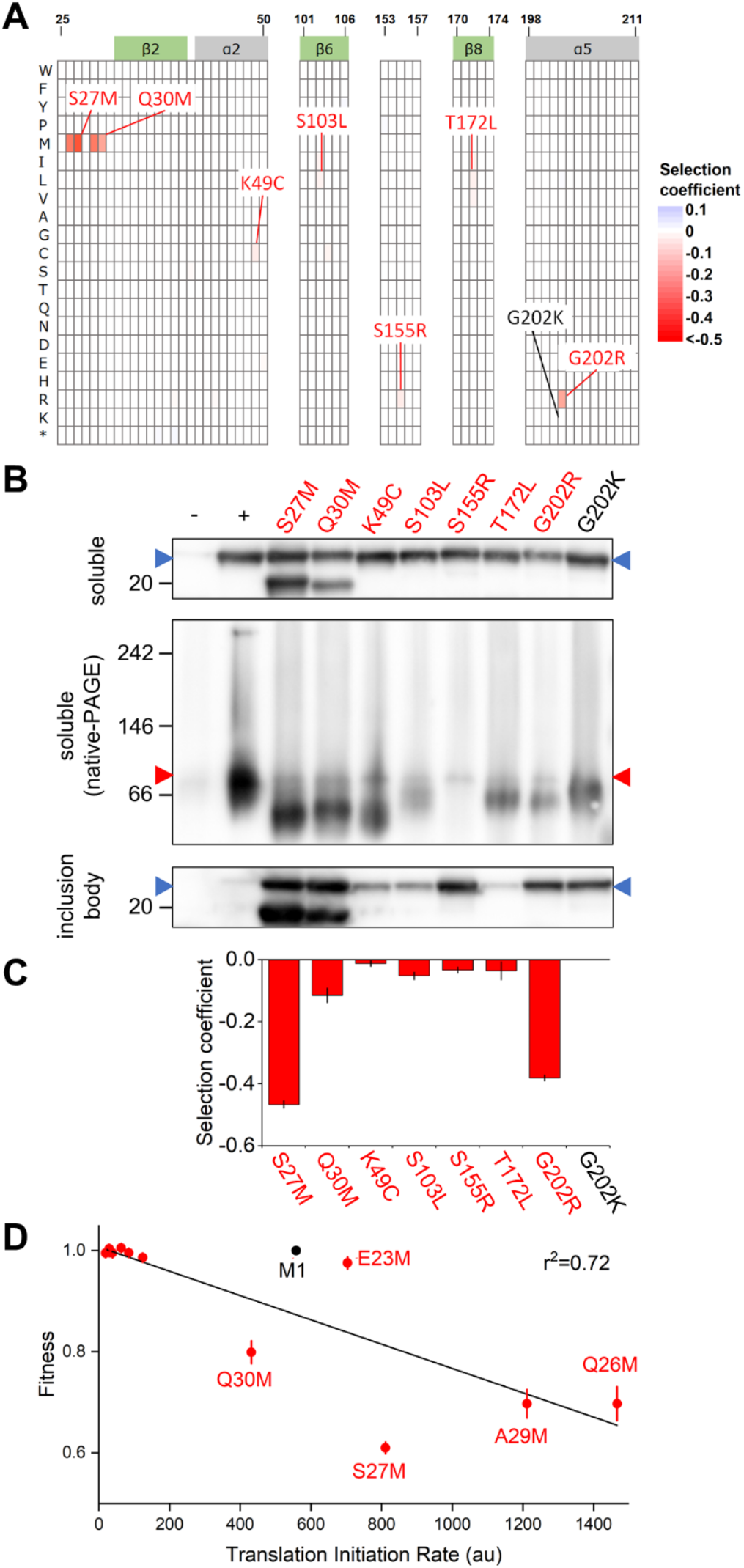
CAT-I mutations’ effect on fitness and protein fate. (**A**) Subsections of the CAT-I collateral fitness landscape showing weighted mean selection coefficients (by DMS) for selected mutations (*P*<0.01 in both replicates). Regions containing β-sheets (green) and α-helices (grey) are labeled along the top. (**B**) Western blots of the total soluble (denaturing and native gels) and inclusion body fractions from cells expressing unmutated CAT-I (–, without IPTG; +, with IPTG) or the indicated CAT-I mutants (deleterious mutations in red and neutral mutations in black). The expected sizes of the monomer (blue arrowhead) and trimer (red arrowhead) are shown. Full western blots including the second biological replicate are provided as **supplementary fig. S11**, Supplementary Material online. (**C**) Selection coefficients were determined from monoculture growth experiments by measuring the mean growth rate over six hours of growth post-induction. Error bars represent 99% confidence intervals (*n*=3). (**D**) Experimental weighted mean fitness values plotted as a function of predicted translation initiation rates at residues 22-34 when the position is mutated to methionine. M1 represents the wild-type start codon and is excluded from the fit. A table of the predicted translation initiation rates and corresponding collateral fitness values for these mutations is included as **supplementary table S1**, Supplementary Material online.

The *aadB* gene was more susceptible to deleterious collateral fitness effects than *NDM-1* or *CAT*-I. We found 3.76% (120/3190) of missense mutations in *aadB* resulted in measurable deleterious collateral fitness effects in both replica experiments (*P*<0.01, *Z*-test) (**fig. 4A**). The fitness effects of deleterious mutations were often small in magnitude (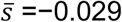,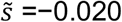 for all deleterious mutations *P*<0.01). We did not find evidence of significant collateral fitness effects for mutations at active site residues R40, H42, D44, D46, and D86 (Cox et al. 2015). We constructed seven mutations in *aadB* chosen as a broad representation of mutations with deleterious effects but focused more on those with large effects. Of these mutations, the three mutations to methionine failed to show a significant fitness effect in monoculture growth, but the other four mutations did (**fig 4C**, **supplementary fig. S5C**, Supplementary Material online).

**Fig. 4.**
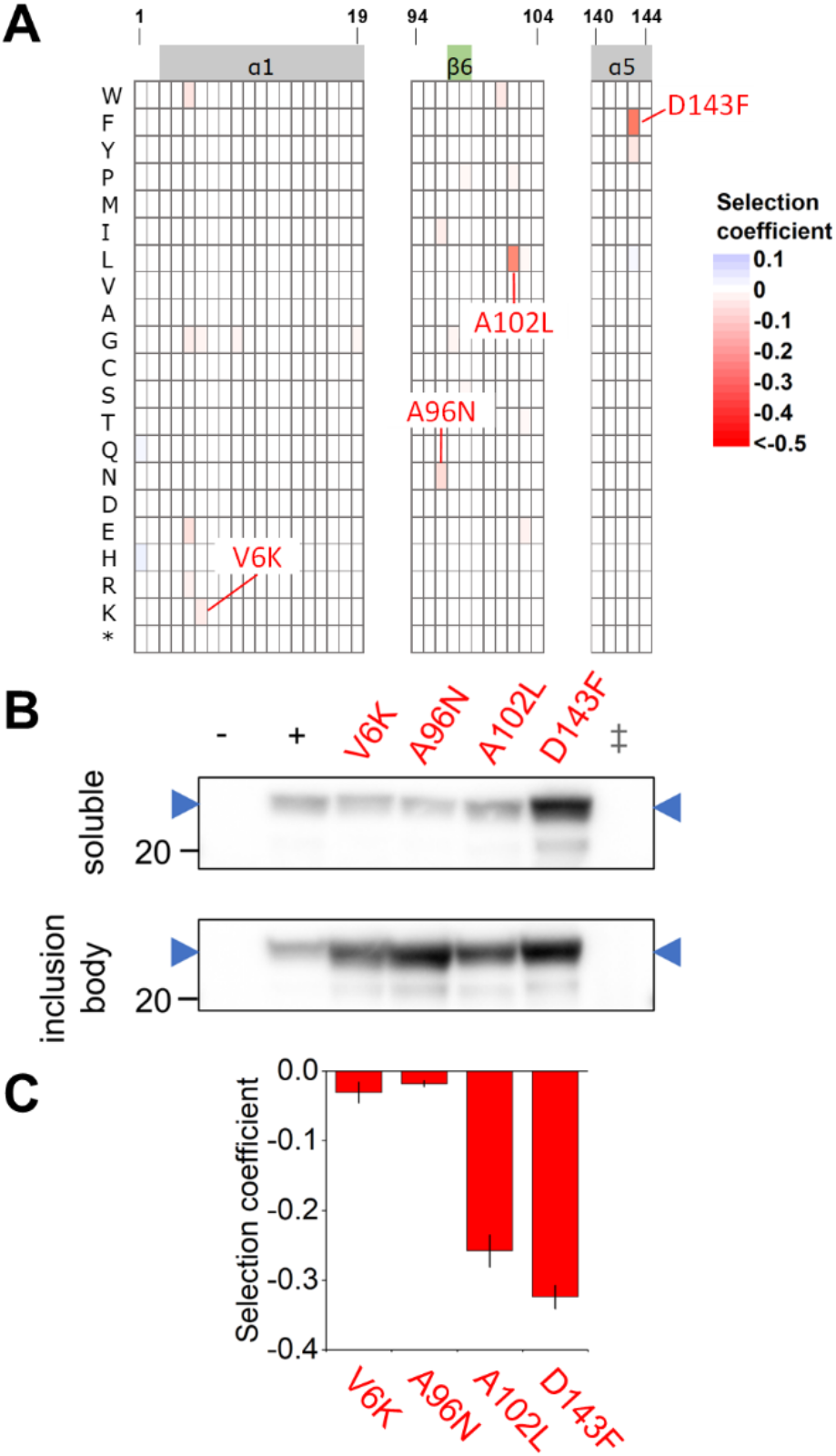
AadB mutations’ effect on fitness and protein fate. (**A**) Subsection of the AadB collateral fitness landscape showing weighted mean selection coefficients (by DMS) for selected mutations (*P*<0.01 in both replicates). Regions containing β-sheets (green) and α-helices (grey) are labeled along the top. (**B**) Western blots of the total soluble and inclusion body fractions from cells expressing unmutated AadB (–, without IPTG; +, with IPTG) or the indicated AadB mutants (deleterious mutations in red and neutral mutations in black). Samples from cells expressing CAT-I (‡) were included to identify the bands of cross-reacting *E. coli* proteins. The blue arrowhead indicates the expected position of AadB. Full western blots including the second biological replicate are provided as **supplementary fig. S12**, Supplementary Material online. (**C**) Selection coefficients were determined from monoculture growth experiments by measuring the mean growth rate over six hours of growth post-induction. Error bars represent 99% confidence intervals (*n*=3).

### Deleterious mutations often cause protein aggregation, but protein aggregation is neither necessary nor sufficient to cause deleterious fitness effects

We next examined how the deleterious and neutral mutations affected the fate of the NDM-1, CAT-I, and AadB proteins using cell fractionation experiments couple with SDS-PAGE and Western blots. In our previous study of TEM-1 mutations with deleterious collateral fitness effects (Mehlhoff et al. 2020), all tested deleterious mutations caused aggregation and signal sequence mutations, in addition, caused improper signal sequence cleavage. Here, our results with NDM-1, CAT-I, and AadB offer a more complicated picture of the relationship between deleterious fitness effects and protein processing/aggregation.

The five most deleterious mutations in *NDM-1* were at residues G25 and C26, positions that are likely to affect export and subsequent lipidation of NDM-1. Like other β-lactamases such as TEM-1, NDM-1 is exported through the Sec export pathway (González et al. 2016; Cheng et al. 2018). However, NDM-1 has an atypical signal peptide and is inefficiently secreted (Zalucki et al. 2020). A canonical Sec signal peptide contains a positively charged N region, a hydrophobic core region, and a neutral C-terminal region that contains the cleavage site immediately after a consensus A–X–A motif (Auclair et al. 2012). NDM-1 has the sequence A-L-A-A-A starting at residue 16, suggesting possible cleavage by signal peptidase I after residue 18 or 20. Studies on NDM-1 with a C-terminal His-tag have identified cleavage after positions 15, 17, 18, and 35 (Thomas et al. 2011) and 21 (Zalucki et al. 2020). However, NDM-1 contains a canonical LSGC lipobox motif at the C-terminus of the signal sequence (residues 23-26), a site that would be cleaved by lipoprotein type II signal peptidase (LspA) after residue 25 (**fig. 2A**). Studies of NDM-1 with a C-terminal Strep-tag found such cleavage after position 25 (González et al. 2016; Prunotto et al. 2020). Thus, the consensus is that NDM-1 is a lipoprotein that anchors to the inner leaflet of the outer membrane in Gram-negative bacteria through C26 after cleavage by LspA (González et al. 2016; Prunotto et al. 2020). Both lipidation and a native affinity of NDM-1 toward the membrane allow the protein to anchor to the inner leaflet of the outer membrane in gram-negative bacteria. NDM-1 anchors in a stable, defined conformation with an exposed active site (Prunotto et al. 2020).

For the wild-type protein, NDM-1 appeared in the total soluble, periplasmic, and inclusion body fractions, but only a small amount appeared in the membrane fraction (**fig. 2B**). The size of the wild-type protein in all fractions suggests that it is the mature protein without a signal sequence. G25 is in the LSGC lipobox and immediately precedes the LspA cleavage site. Mutations G25C, G25S, and G25T, but not any other mutation at G25, caused large deleterious effects (**fig. 2AC**). G25C and G25S did not appreciably alter the amount or size of NDM-1 in the soluble or inclusion body fractions, and the species was detected in the periplasmic fraction (**fig. 2B)**. In contrast, the G25T mutation led to a drastic reduction in levels of mature NDM-1 in the soluble and insoluble fractions and the appearance of a higher molecular weight band in both fractions, suggesting G25T disrupts regular cleavage of the signal sequence. No NDM-1 species with the G25T mutation could be detected in the periplasmic or membrane fractions.

At position 26 we characterized the highly deleterious mutations C26S and C26D and compared to the neutral mutations C26L and C26A (**fig. 2B**). C26A has been shown to result in soluble NDM-1 accumulating in the periplasm because it removes the lipidation site, thereby preventing membrane anchoring (González et al. 2016). This suggests mutations at C26 can disrupt signal sequence cleavage with a difference in association with the membrane. All four mutations reduced mature NDM-1 in the soluble, inclusion body, and membrane fractions to an undetectable level (**fig. 2B**). C26S caused the NDM-1 precursor to accumulate in the membrane fraction while the C26D and C26A mutations caused the precursor to accumulate in the periplasm. Although these results indicate C26S and C26D decreases precursor processing, the fact that the neutral mutation C26A does so as well suggests impaired removal of the signal sequence may not be the cause of the deleterious collateral fitness effects of C26S and C26D.

We also characterized the neutral P28A mutation because it is the sole mutation differentiating the naturally occurring NDM-2 variant from NDM-1. The mutation has been noted to result in a more canonical signal peptidase I cleavage site that improves secretion and leads to soluble, non-lipidated protein in the periplasm (Bahr et al. 2018; Zalucki et al. 2020). In contrast, we find the P28A mutation led to a reduction of NDM-1 in the soluble and periplasmic fraction, an increase of NDM-1 in the inclusion body fraction, and no evidence of a reduction in the small amount of NDM-1 in the membrane fraction. Despite the increase in aggregation, P28A did not cause deleterious collateral fitness effects (**fig. 2B**). In conclusion, neither impaired signal sequence cleavage nor increased aggregation is sufficient or necessary to cause deleterious collateral fitness effects in NDM-1.

All deleterious CAT-I mutations produced wild-type levels of soluble protein, but unlike wild-type, most led to accumulation of CAT-I in the inclusion body fraction (**fig. 3B**). However, G202K, a neutral mutation tested to compare with the highly deleterious G202R, also caused aggregation despite its lack of a fitness effect. Aggregation of G202 mutants is consistent with the observed importance of a full-length α5-helix (residues 198-211) for correct folding and stabilization of CAT-I (Van der Schueren et al. 1996). Similarly, non-hydrophobic mutations at C212 have been reported to cause inclusion body formation (Van der Schueren et al. 1996), yet such substitutions did not cause deleterious collateral fitness effects (the weighted mean *s* was −0.004 with a 99% confidence interval of 0.015). Thus, aggregation of CAT-I per se is not sufficient for deleterious collateral fitness effects. If aggregation of some mutants causes deleterious effects, then it must be because the specific nature of the aggregation is deleterious. For example, perhaps the deleterious mutant aggregates with another protein, causing a deleterious fitness effect, but the neutral mutant aggregates only with itself.

Methionine substitutions at Q26, S27, A29, and Q30 in CAT-I led to large-magnitude deleterious collateral fitness effects in the DMS data with selection coefficients in the range of −0.20 to −0.39. We chose to study S27M and Q30Mas examples of these mutants. Both mutations resulted in truncated and full-length versions of CAT-I in the soluble and inclusion body fractions (**fig. 3B**). We hypothesized that the truncated version originated from the mutation causing translation initiation at these sites and that the truncated protein was the cause of the deleterious effect. To test this hypothesis, we predicted the mRNA’s translation initiation rate at positions 22-34 when each codon was replaced with the start codon ATG (Reis and Salis 2020). Consistent with our hypothesis, the fitness effect for methionine substitutions correlated with predicted mRNA translation initiation rates (*r*^2^ = 0.72) (**fig. 3D**)

One deleterious mutation occurred at a residue important for trimer formation. S155 is involved in formation of the trimeric core in the CAT-I homotrimer (Goodale et al. 2020) although the residue is weakly conserved and substituted by glycine in certain CAT homologs (Biswas et al. 2012). S155R was the only missense mutation at this position with a reproducible, deleterious effect (*s* =−0.034 ± 0.01). It led to CAT-I aggregates in the inclusion body fraction and almost complete loss of trimer formation as monitored by native gel electrophoresis despite the monomer being present in the soluble fraction (**fig. 3B**). Several deleterious mutations (but not the G202K neutral mutation) led to the putative trimeric species running slightly faster during native gel electrophoresis. This observation suggests that these mutations altered the nature of the trimer, though the S27M and Q30M mutants’ faster migration can be explained by the presence of N-terminally truncated species.

For AadB, we characterized the effect on protein fate of the confirmed deleterious mutations V6K, A96N, A102L, and D143F. Like NDM-1, wild-type AadB was found both in the soluble and inclusion body fractions (**fig. 4B**). Only D143F altered the levels of soluble AadB, surprisingly causing a substantial increase in soluble protein. All four mutations caused an increase in AadB aggregates, although not at levels proportional to the size of their respective fitness effects.

### Select missense mutations can cause deleterious collateral fitness effects without affecting physiological function

We examined the relationship between primary and collateral fitness effects for several mutants using a minimum inhibitory concentration (MIC) assay as a proxy for primary fitness measurements (**fig. 5A-C**). For many mutants, collateral fitness effects roughly correlated with primary fitness effects. Cells expressing mutants with little or no collateral fitness cost tended to retain near-full antibiotic resistance, and those mutants in *NDM-1* and *CAT-I* with the most severe collateral fitness cost completely lost the ability to confer antibiotic resistance. Notable exceptions were the S155R and T172L mutations in *CAT-I*, which had much larger primary fitness effects. This differential impact on primary and collateral fitness effects is expected, even common, as it is easy to imagine how a mutation can affect catalytic activity without causing collateral fitness effects. For example, mutations at residues in and near the active site of NDM-1 often lead to a substantial loss in resistance (Sun et al. 2018), but almost all lacked a measurable collateral fitness cost in our study. The inverse case, in which mutations with substantial collateral fitness effects had little or no primary fitness effects, is less intuitive but was observed with *TEM-1* (Mehlhoff et al. 2020). For the genes studied here, mutations G25T in *NDM-1* and D143F in *aadB* had a collateral fitness cost of 23% and 32%, respectively, yet retained the wild-type ability to confer antibiotic resistance as measured by their MIC. Because G25T prevents mature protein formation (**fig. 2B**), this means that preNDM-1 with G25T possesses catalytic activity. D143F results in an increase in soluble protein (**fig. 4B**), which may help the mutant retain the ability to confer full resistance.

**Fig. 5.**
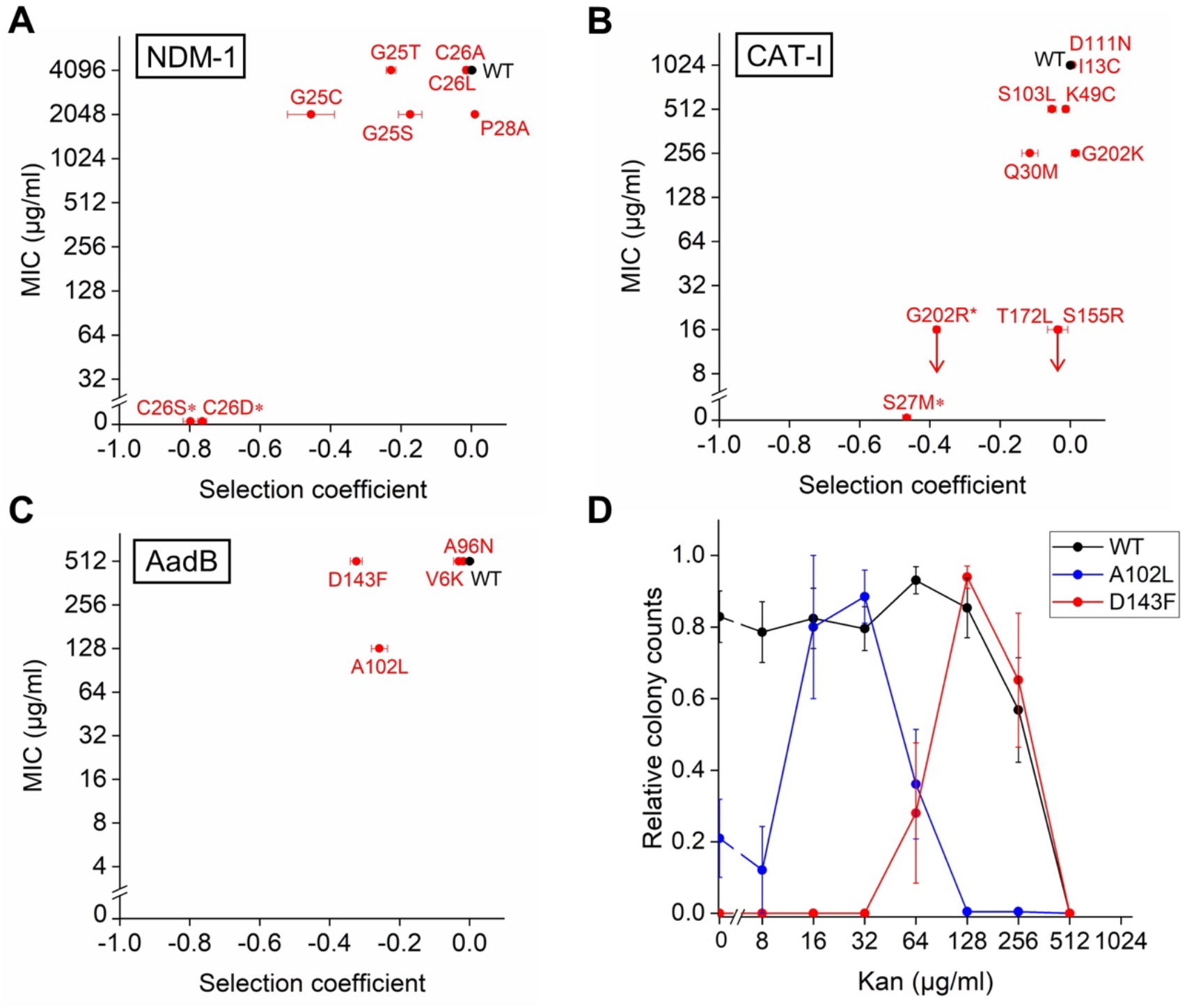
Comparison of primary and collateral fitness effects. We utilized a minimum inhibitory concentration (MIC) assay in which cells expressing wild-type or mutants of the antibiotic resistance genes were grown overnight with IPTG inducer and plated on agar plates containing the corresponding antibiotic. Plots show the relationship between the collateral fitness effects and the median MIC values for cells expressing (**A**) *NDM-1* variants plated on ampicillin-containing plates, (**B**) *CAT-I* variants plated on chloramphenicol-containing plates, and (**C**) *aadB* variants plated on kanamycin-containing plates as a function of collateral fitness effects as measured by monoculture growth. Error bars represent the 99% confidence interval. Red arrows indicate a MIC of <16 μg/ml for G202R, T172L, and S155R. Variants that did not form colonies on a plate lacking antibiotics in at least one replicate of the assay are marked with an (*). For *CAT-I*, the points for S155R and T172L overlap. (**D**) Mean relative colony counts in the MIC assays for the wild-type *aadB* and A102L and D143F mutants. Error bars are standard error of the mean (n=5). Plots of mean relative colony counts for all MIC assay samples are included as **supplementary fig. S13**, Supplementary Material online. A complete table of MIC values is provided as **supplementary table S2**, Supplementary Material online with colony counts from the assay available as **supplementary data S5**, Supplementary Material online.

### Two AadB mutations cause an unusual antibiotic resistance phenotype

The two *aadB* mutations causing the largest deleterious collateral fitness effects, A102L and D143F, also caused an unusual and unexpected antibiotic resistance phenotype: band-pass resistance behavior (**fig. 5D**). Cells with the D143F mutation grew at kanamycin concentrations as high as 256 μg/ml, but these cells were unable to grow at any concentration of kanamycin ≤ 32 μg/ml including, remarkably, the absence of kanamycin. Similarly, cells with A102L grew on plates containing 16 or 32 μg/ml kanamycin but formed fewer colonies on plates with either lower or higher levels of antibiotic. Such band-pass behavior in antibiotic resistance has been engineered before using a synthetic gene circuit (Sohka et al. 2009). In that case, band-pass behavior was achieved by requiring that the ratio of catalytic activity to antibiotic concentration is high enough to degrade enough of the antibiotic so that the cells can grow, but not too high to reduce the antibiotic concentration below the threshold needed to induce expression of an essential gene. Perhaps the presence of sub-lethal amounts of kanamycin induces a cell response that ameliorates the deleterious collateral fitness effects caused by D143F. Alternatively, having an abundance of the substrate kanamycin may serve to stabilize AadB’s folded structure through the energy of binding, preventing aggregation or misinteractions and ameliorating the deleterious collateral fitness effects present at low or no substrate.

### Select stress-response pathways are activated in cells expressing deleterious mutants

We utilized RNA sequencing (RNA-Seq) on a subset of mutations in *CAT-I*, *NDM-1*, and *aadB* to examine how the cell’s transcriptome responds to mutations causing deleterious collateral fitness effects. We calculated the fold-difference in expression levels of genes from cells expressing a mutant relative to the expression levels of those genes in cells expressing the wild-type antibiotic resistance protein. We identified genes and select pathways in which mutations caused a more than two-fold change in expression with *P*<0.001 statistical significance for multiple interconnected genes or for stress pathways. We included a control consisting of a culture of cells that lacked the antibiotic resistance gene but contained the IPTG expression inducer. A comparison of this control to a culture containing the wild-type *aadB* gene showed expression of AadB had minimal effects on the transcriptome (**supplementary fig. S2C**, Supplementary Material online). For cells containing any of the three antibiotic genes, the addition of the IPTG inducer did not cause significant changes in the transcriptome except for upregulation of the *lac* operon, as expected (**supplementary fig. S2D-F**, Supplementary Material online).

We observed that mutations lacking a significant fitness effect (G202K in *CAT-I* and P28A in *NDM-1*) or mutations that were only slightly deleterious (S155R in *CAT-I*) had much fewer and smaller-magnitude effects on the transcriptome than deleterious mutations (**supplementary fig. S14**, Supplementary Material online). However, the selection coefficient was a poor predictor of the frequency of genes with perturbed expression (**supplementary fig. S14D**, Supplementary Material online). For example, even though G202R in *CAT-I* (*s* = −0.38) was slightly more deleterious than the A102L and D143F mutations in *aadB*, the *aadB* mutations caused changes in expression eight times more frequently than G202R (5.3% for G202R vs. 37.8% and 42.6% for A102L and D143F, respectively). In fact, the *aadB* mutations so perturbed the transcriptome that it was difficult to identify changes to focus on. We instead focus our discussion on the effects of *CAT-I* and *NDM-1* mutations in comparison with our previous results with *TEM-1*.

We previously showed that deleterious mutations in *TEM-1* strongly activated two periplasmic stress pathways: the phage shock protein (Psp) response and the regulator of colanic acid capsule synthesis (Rcs) pathway, with the exception that most deleterious signal sequence mutations did not activate the Rcs response despite causing some of the largest collateral fitness effects (Mehlhoff et al. 2020). The Psp response is believed to be induced by inner membrane stresses and may work to mitigate secretin toxicity or problems stemming from increased inner membrane permeability (Flores-Kim and Darwin 2016). In contrast, the Rcs pathway is commonly induced by outer membrane or peptidoglycan layer stresses along with disruption of periplasm functions. It triggers positive regulation of capsular polysaccharides for biofilm formation and negative regulation of motility (Wall et al. 2018).

Here, we find that deleterious mutations in NDM-1, which are at the proposed signal sequence cleavage and lipidation sites, activated both pathways (**fig. 6AB**). Like TEM-1, NDM-1 contains a signal sequence and requires export to the periplasm for folding and function. Deleterious mutations surrounding the proposed signal peptidase cleavage site in NDM-1 caused 6- to 142-fold upregulation of genes corresponding to the Psp stress response (*P*<10^−14^). Because G25T, C26S, and C26D completely prevented mature protein formation, they may disrupt typical export across the inner membrane or signal sequence cleavage, causing deleterious collateral fitness effects leading to the activation of the Psp pathway. In agreement with that hypothesis, no deleterious mutation in the cytoplasmic proteins CAT-I or AadB activated the Psp pathway (**fig. 6A**). However, G25C and G25S did not alter the amount of NDM-1 that was correctly exported and cleaved, suggesting these mutations must activate the Psp stress pathway via a different mechanism.

**Fig. 6.**
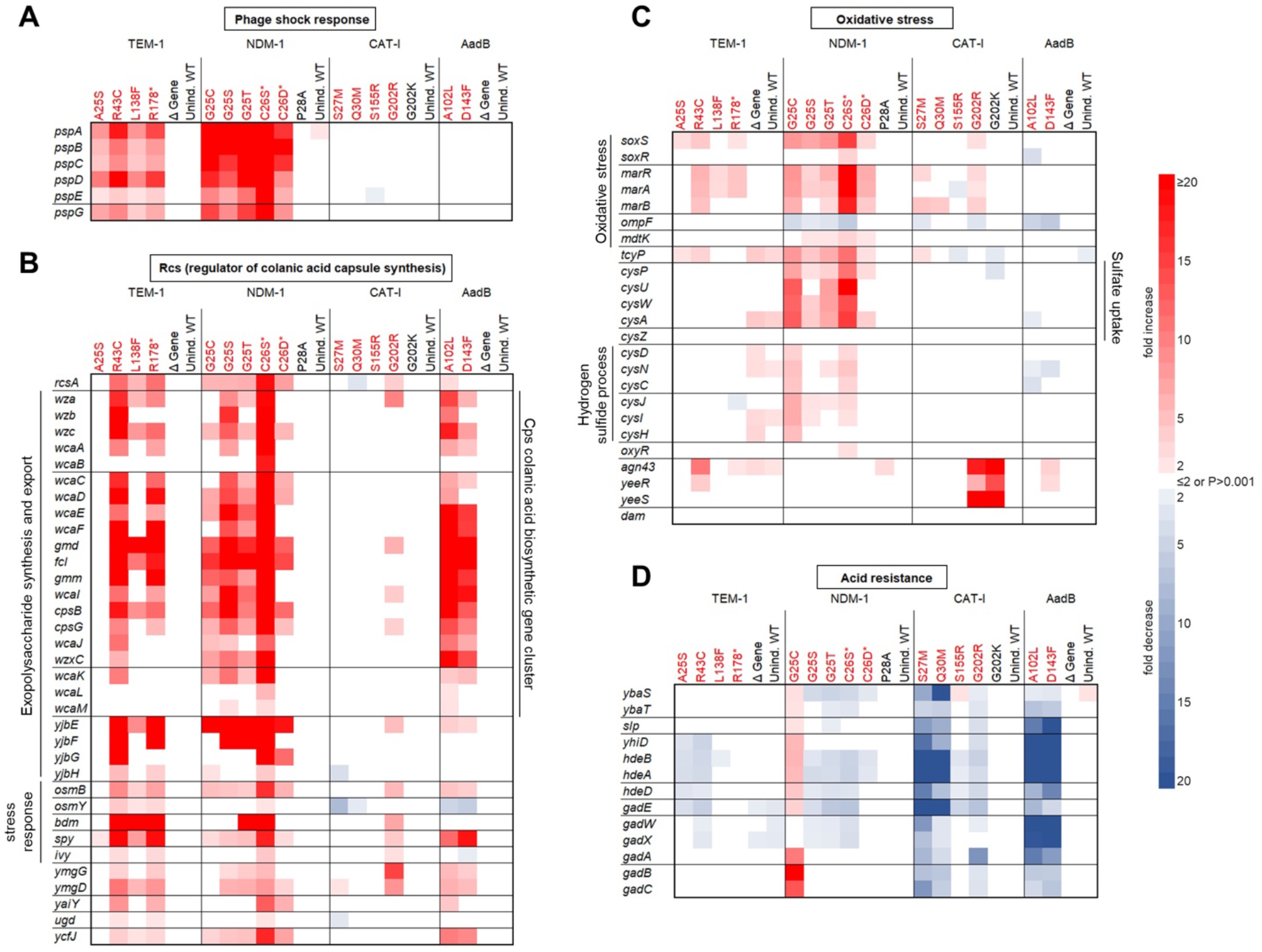
Effects of mutations with deleterious collateral fitness effects on the*E. coli* transcriptome. We collected exponentially growing cells after approximately 330 minutes of induced growth and performed RNA-Seq experiments to monitor expression levels of genes throughout the transcriptome. The heat maps show select genes in the (**A**) phage shock protein (Psp) response (**B**) regulator of colanic acid capsule synthesis (Rcs) pathway (**C**) oxidative stress responses, and the (**D**) GAD acid resistance response that differed by greater than twofold expression (*P*<0.001, Z-test) relative to cells expressing the wild-type antibiotic resistance genes. The color of the mutation indicates whether it is deleterious (red) or neutral (black). Control samples include ones in which wildtype gene expression was not induced (Unind. WT) and induced ones which lack the gene (Δ Gene). RNA-Seq data from representative mutations in *TEM-1* is from Mehlhoff *et al*. (Mehlhoff et al. 2020). Additional observations of gene expression are provided as **supplementary text**and **fig. S16**, Supplementary Material online. Tabulated RNA-Seq data is provided as **supplementary data S4**, Supplementary Material online.

Signal sequence mutations in NDM-1 but not TEM-1 activated the Rcs pathway, a difference that may result from NDM-1 being a lipoprotein that anchors to the outer membrane while TEM-1 is not. However, certain mutations in both cytoplasmic proteins caused upregulation of the Rcs pathway (A102L and D143F in *aadB* and, to a lesser extent, G202R in *CAT-I*) (**fig. 6B**). This suggests that mutational consequences in the cytoplasm can cascade to activate the Rcs pathway.

Like *TEM-1*, deleterious mutations in *NDM-1* did not cause activation of two other periplasmic stress responses (σ^E^ and BaeR) but did show mild activation of the Cpx envelope response (**supplementary fig. S15A**, Supplementary Material online). The Cpx envelope stress-response is thought to help retain inner membrane integrity and is stimulated by inner membrane stress as well as misfolded inner membrane or periplasmic proteins (Raivio 2014; Grabowicz and Silhavy 2017). The S27M and Q30M methionine substitutions in *CAT-I* along with the highly deleterious *aadB* mutations caused a slight upregulation of genes in the σ^H^-mediated heat shock response (**supplementary fig. S15B**, Supplementary Material online).

All five highly deleterious *NDM-1* mutations caused changes in gene expression indicative of periplasmic oxidative stress (**fig. 6C**). First, the expression of *soxS* and its paralog, *marA*, were elevated 3- to 15-fold (*P*<10^−15^) and 4- to 27-fold (*P*<10^−11^), respectively. Both genes encode transcriptional activators that regulate an overlapping set of genes, with SoxS helping in the removal of superoxide and MarA helping in resistance to antibiotics (Holden and Webber 2020). The increased expression of these activators did not result in changes in expression of most of their target genes except for a 3- to 6-fold reduction in *ompF* (*P*<10^−15^), which encodes an outer membrane porin and a 2- to 3-fold increase in *mdtK* (*P*<10^−4^), which encodes a multidrug efflux transporter. The second indicator of oxidative stress was an upregulation of the cysteine/cystine shuttle system, which plays an important role in oxidative stress tolerance by providing reducing equivalents to the periplasm (Mironov et al. 2020). In the system, hydrogen peroxide in the periplasm is removed in the oxidation of cysteine to cystine. Cystine is imported to the cytoplasm via TcyP (3- to 12-fold increase in expression, *P*<10^−8^). In the cytoplasm, cystine is reduced to cysteine so it can be shuttled back to the periplasm to remove more hydrogen peroxide. Consistent with a need for more cysteine, which is synthesized from serine and hydrogen sulfide, most genes in the *cysDNC and cysJIH* pathway for converting sulfate to hydrogen sulfide and the *cysPUWA* operon for importing sulfate/thiosulfate were upregulated in the NDM-1 mutants (**fig 6C**). The third indicator of oxidative stress was the upregulation of genes for the synthesis of enterobactin, which aside from its role in iron uptake also plays a role in the oxidative stress response (**supplementary fig. S15C**, Supplementary Material online) (Peralta et al. 2016). The cytoplasmic oxidative stress regulator *oxyR* (**fig 6C**) and the genes OxyR activates when the cytoplasm experiences oxidative stress due to hydrogen peroxide (e.g. *sufABCDSE, katG, poxB, hemH, mntH*, and *grxA*) (Roth et al. 2022)did not show elevated expression.

### A mutation in CAT-I lacking a collateral fitness effect can nonetheless cause specific changes in gene expression

Interestingly, we observed that a mutation without a collateral fitness effect could cause a precise and large change in gene expression. Both G202R and G202K cause CAT-I aggregation (**fig. 3B**) yet only G202R causes a collateral fitness effect (*s* = −0.38 ± 0.01 and 0.01 ± 0.01, respectively). Surprisingly, both mutations (but not the other three CAT-I mutations) caused a dramatic increase in *agn43* expression (18- and 29-fold, respectively, *P*<10^−15^), a gene that encodes two outer membrane proteins (**fig. 6C**). Both mutations also caused similar expression increases in *yeeRS* immediately downstream (**fig. 6C**). Expression of *agn43* is blocked by binding of OxyR to the *agn43* promoter (independent of OxyR’s oxidative state), but *dam* methylation of the *agn43* promoter blocks OxyR binding. Competition between Dam and OxyR determine *agn43* expression (Kaminska and Van Der Woude 2010). Because neither *dam* nor *oxyR* expression was altered by the mutations (**fig. 6C**), the mutations may cause CAT-I to interfere with OxyR binding, causing high *agn43* expression. This might occur if the mutation causes CAT-I to bind OxyR and sterically block OxyR’s interaction with the agn43 promoter or if the mutation causes co-aggregation of CAT-I and OxyR, depleting the amount of OxyR that can bind the agn43 promoter. Ag43 has been linked to the oxidative stress response and biofilm formation (Laganenka et al. 2016).

### Mutations at G25 in NDM-1 cause opposite changes in gene expression

We observed one instance in which two mutations at the same position caused opposite changes in expression of a stress response pathway. Many deleterious mutations across all four antibiotic resistance genes caused a decrease in expression of *gadE*, *gadX*, and *gadW* (**fig. 6D**), as well as repression of genes activated by GadX/GadW (all the genes listed above *gadE* in **fig. 6D**) as part of the glutamate-dependent acid resistance response (Tucker et al. 2003), which helps restores pH homeostasis in acidic environments. However, the G25C mutation in *NDM-1* caused activation of this acid resistance response with a 10- to 23-fold increase in expression of *gadA* and *gadBC* (*P*<10^−15^), in sharp contrast to G25S and G25T. GadA and GadB catalyze the H^+^-consuming conversion of glutamic acid to γ-aminobutyrate (De Biase et al. 1999). Neither G25C nor G25S alter the fate of NDM-1 (**fig. 2B**) and otherwise produce very similar changes to the transcriptome (**fig. 6** and **supplementary figs. S14-S16**, Supplementary Material online). Because Cys and Ser only differ by one atom (sulfur vs. oxygen), this suggests that the reactivity of a cysteine in the G25C mutation causes the differential activation of the glutamate-dependent acid resistance response, though differences in protein dynamics could also be the source of the difference.

### S27M and Q30M mutations in CAT-I cause loss of the F’ episome

Our cells contain a large, single-copy F’ episome in addition to the pSKunk-3 plasmid encoding the tested genes. We chose a strain with this F’ episome because it encodes an extrachromosomal copy of the *lacI* repressor gene, which better represses the expression of our target antibiotic resistance genes before the growth competition. This helps us maintain deleterious mutants in the library during the library creation and propagation steps before the growth competition experiment, which is initiated by inducing expression of our target genes. Under typical circumstances, the F’ episome does not require a maintenance antibiotic to be stably propagated to daughter cells.

We observed the S27M and Q30M methionine substitutions in *CAT-I* caused a dramatic decrease in expression of conjugation genes on the F’ episome (**supplementary fig. S15D**, Supplementary Material online). In fact, both mutations caused a drastic reduction in counts for almost all episomal genes, often reducing the counts to zero (**fig. 7A**). In addition, *lacI* expression decreased 5- to 11-fold. Suspecting the mutations might be causing loss of the episome, we used a purification method designed for isolation of large DNA molecules, performing the extraction after 10 generations of induced monoculture growth for the three most deleterious mutations in CAT-I, including S27M and Q30M. PCR amplification of the *traJ* and *traI* episomal genes from this fraction revealed the reduction or complete loss of the episome from cells expressing S27M and Q30M but not for G202R (**fig. 7B,C**). TraJ is the master regulator of *tra* operon transcription while TraI is a relaxase responsible for the beginning of the conjugation process. The most straightforward potential mechanism is that the CAT-I protein fragments resulting from the S27M and Q30M mutations interfere with episome replication, maintenance, or segregation during cell division, causing a reduction in episome propagation. Alternatively, the fragments might interact with an episomal protein such that the presence of the episome becomes toxic, resulting in selection against episome-containing cells.

**Fig. 7.**
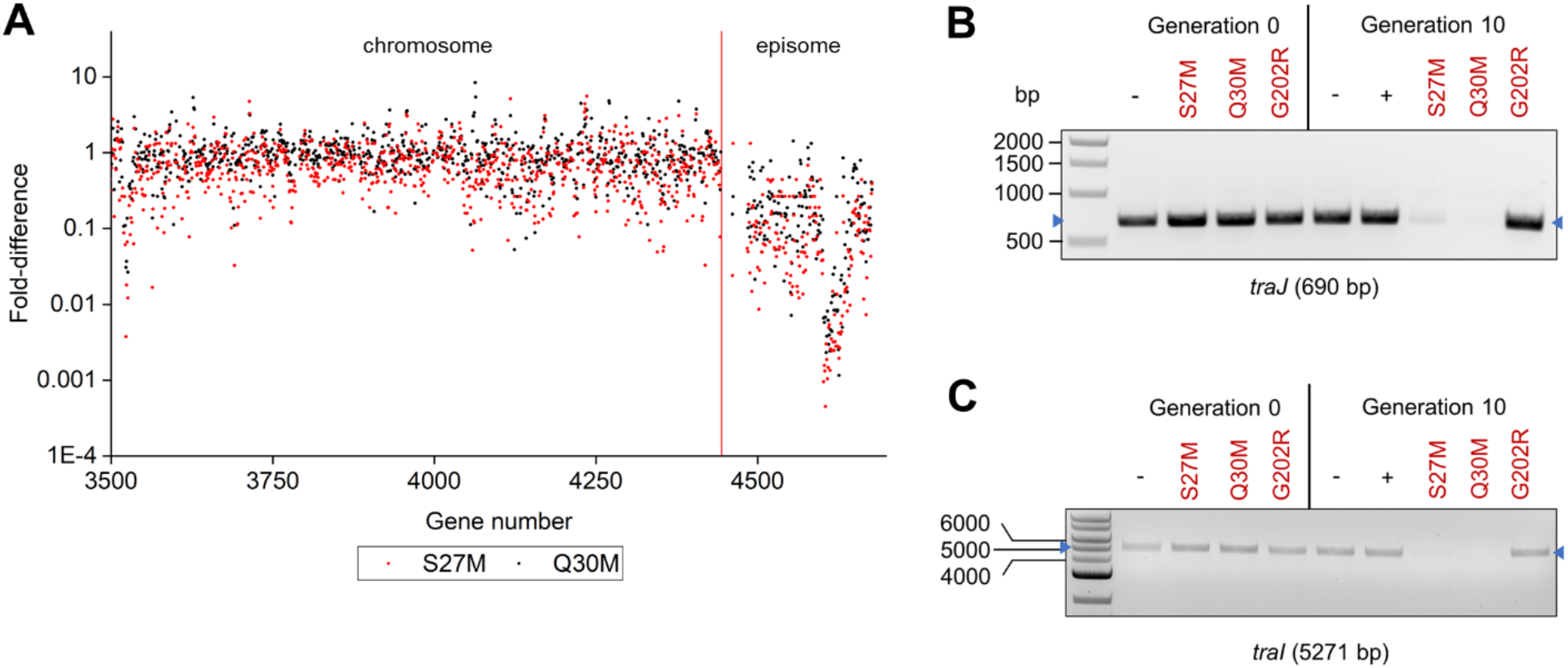
Loss of episome corresponding to deleterious methionine substitutions in CAT-I. RNA-Seq revealed the S27M and Q30M methionine substitutions in *CAT-I* led to (**A**) reduced expression of episomal genes after 10 generations of induced growth. All genes to the right of the red line are on the F’ episome. PCR amplification of the (**B**) *traJ* and (**C**) *traI* episomal genes illustrate the loss of these genes in cells expressing the S27M or Q30M mutants of CAT-I. The G202R CAT-I mutation represents a mutation causing large-magnitude deleterious effects (like S27M and Q30M) but no change in *traJ* and *traI* expression.

## Discussion

Our study shows that genes have different propensities to exhibit collateral fitness effects. What are the origins of these differences? One possible explanation is differences in a protein’s propensity for aggregation/misfolding. All the deleterious mutants studied in *TEM-1* and *aadB* increased aggregation, and these two genes had the highest frequency of mutations with deleterious fitness effects. This correspondence is consistent with the hypothesis that a protein’s propensity for aggregation predicts the frequency of deleterious collateral fitness effects. However, our study shows that not all aggregation causes deleterious fitness effects. Unmutated NDM-1 aggregates, and the only NDM-1 mutation studied that increased aggregation (P28A) had neutral fitness effects (**fig. 2**). G202R and G202K in CAT-I cause approximately the same amounts of aggregation, but only G202R was deleterious (**fig. 3**). For mutants of CAT-I and AadB, there was little correlation between the magnitude of the deleterious fitness effect and the amount of aggregation (**figs. 3** and **4**). The aggregation we observed cannot be deleterious to fitness due to a resource cost because then the magnitude of the effect and the amount of aggregate should positively correlate, and such a resource cost would be very small relative to the fitness effects measured here. Instead, for aggregation/misfolding to be deleterious for fitness, the aggregate/misfolded protein must misinteract with a cellular component or process in a manner that is deleterious for fitness. For example, perhaps G202R aggregates in a toxic way but G202K does not.

This perspective is important when evaluating evidence addressing the role aggregation/misfolding may play in the E-R anticorrelation. Agozzino and Dill’s theoretical treatment found that protein evolutionary rates are faster for less stable proteins (Agozzino and Dill 2018). In contrast, Usmanova et al. found that experimental measures of protein stability and melting temperature do not correlate with evolutionary rate (Usmanova et al. 2021). The authors interpreted this lack of correlation as undermining the argument that misfolding plays a major role in the E-R anticorrelation. However, their interpretation is based on the premise that less stable proteins will be more likely to aggregate upon mutation because their stability buffer is small. However, most proteins are marginally stable (Taverna and Goldstein 2002), virtually all proteins have folding intermediates, and folding and misfolding in the cell is a kinetic competition (Jahn and Radford 2008). Although destabilization via mutation will increase the population of partially folded conformations, the key for aggregation is how the mutations affect the nature of those intermediates and the barriers that must be overcome for the folding protein to progress to aggregation. Thus, what may be important is not just whether the mutation can sufficiently compromise the global stability of the native state, but how the mutation affects the kinetics of aggregate formation versus native folding and, most certainly, whether the aggregates interact with other cellular components to cause deleterious fitness effects.

An additional potential reason for the differences between genes in the frequency of collateral fitness effects is that chaperones and stress-response pathways likely have different abilities to alleviate the deleterious effects of mutations for the different proteins. For example, the cell’s chaperones and stress responses may be more capable of buffering the negative effects caused by CAT-I and NDM-1 mutations than those caused by TEM-1 and AadB mutations. Consistent with this theory, proteins that interact with chaperones in yeast have been noted to evolve more quickly than proteins that do not (Alvarez-Ponce et al. 2019).

Given the prevalence of mutants causing aggregation, we find it curious that we observed only mild activation of two of the stress-response pathways known to respond to protein aggregation. Instead, the most commonly activated stress responses are two outer-envelope stress pathways: the Psp response induced by mutants of the periplasmic proteins and the Rcs response induced by most mutants in *TEM-1*, *NDM-1*, and *aadB* (and G202R in *CAT-I*). It is unknown whether these stress-response pathways alleviate some of the deleterious fitness effects or whether deleterious fitness effects are brought on by the useful or errant induction of these or other pathways.

We previously proposed that deleterious collateral fitness effects may play a role in protecting against genetic drift toward mutations that would otherwise compromise the protein’s ability to perform its physiological function (Mehlhoff and Ostermeier 2020). Our MIC assays on select mutations with deleterious collateral fitness effects (**fig. 5**) provide only a small snapshot of the overlap of primary and collateral fitness effects in CAT-I, NDM-1, and AadB. Mutations with large deleterious collateral fitness effects also caused deleterious effects in the presence of antibiotic. This co-occurrence of collateral and primary fitness effects, which we also observed in more extensive analysis of *TEM-1* (Mehlhoff et al. 2020), has potential evolutionary benefits under two scenarios. First, in environments in which a protein is expressed but is not under selection for its physiological function, mutations with deleterious collateral fitness effects will be purged. Because these tend to be mutations that also cause primary fitness effects, the population is spared having deleterious mutations should the environment change to one in which the physiological function is relevant for fitness. Similarly, when primary fitness effects are buffered by excess in expression or activity, deleterious collateral fitness effects will tend to prevent the accumulation of mutations that erode this buffer.

Our study provides several examples illustrating how the cell is a complex molecular space that is rich with the potential to interact with mutations in unexpected ways, altering fitness but also causing new phenotypes to emerge. The S27M and Q30M mutations in CAT-I cause loss of the F’ episome – a normally stable extra chromosomal element. In NDM-1, G25C activates but G25S represses the glutamate-dependent acid resistance response. And most strikingly, D143F in AadB makes kanamycin, the antibiotic AadB inactivates, essential for growth. The interactions between mutations and the cell are both an evolutionary constraint and an evolutionary opportunity.

There are several lines of research that would further our understanding of the mechanisms of collateral fitness effects, the factors governing a gene’s susceptibility to them, and the causes and roles of changes in gene expression. One is to use biochemical techniques to identify the biomolecules that interact with or are directly affected by the mutant proteins. Another is to examine how collateral fitness effects are impacted by the environment. For example, protein aggregation is typically lowered by reducing the growth temperature or slowing the growth rate by using a minimal growth media. Do those environmental changes also reduce collateral fitness effects? Specific to NDM-1, we studied the protein in the zinc-rich environment of LB-media. However, evidence suggests that zinc deprivation, and not enzymatic function, is the primary selective pressure acting on the evolution of NDM variants (Bahr et al. 2018). A study of collateral fitness effects in a zinc-limited environment may provide better insight into the collateral fitness effects of mutations under more evolutionarily relevant conditions. A third approach is a genetic one. How do collateral fitness effects depend on the strain of bacteria? Does genetically removing the ability of a cell to activate a stress pathway alter the magnitude of a collateral fitness effect? Does overexpression of chaperones lessen the fitness effects? A fourth approach is to investigate the role of protein expression level. A DMS study conducted in yeast on the green fluorescent protein found at least 42% of the collateral fitness effects caused by single-nucleotide mutations in the protein were dependent on expression level (Wu et al. 2022). They found similar expression-dependent effects in a conditionally essential gene and concluded that increased expression levels in non-essential genes tended to exacerbate deleterious collateral fitness effects. However, their analysis indicated that <6% of the variability in fitness could be explained by misfolding, misinteractions, or translation errors, but that may reflect the accuracy of their measurements and predictions given the small fitness effects they observed (Wu et al. 2022).

## Materials & Methods

### Plasmid construction and general growth conditions

The *TEM-1*, *NDM-1*, *CAT-I*, *aadB*, and *aac(6’)-Im* antibiotic resistance genes were individually placed under control of the IPTG-inducible *tac* promoter on pSKunk1, a minor variant of plasmid pSKunk3 (AddGene plasmid #61531) (Firnberg and Ostermeier 2012). The *CAT-I* gene was amplified from pKD3 (AddGene plasmid #45604). An A to C mutation was made at base pair 219 within the *CAT-I* gene using the QuickChange Lightning Site-Directed Mutagenesis kit (Agilent) to match the native *E. coli CAT-I* sequence. The *NDM-1, aadB*, and *aac(6’)-Im* genes were ordered as gene fragments with adapters from Twist Bioscience. We verified the correct size and antibiotic resistance gene sequence for each plasmid using agarose electrophoresis gels and Sanger sequencing, respectively before transforming the resulting plasmids into electrocompetent NEB 5-alpha LacI^q^ cells, which contain an F’ episome encoding LacI. We produced these electrocompetent cells starting from a single tube of NEB 5-alpha LacI^q^ chemically competent cells. Chemically competent cells were plated on LB-agar containing 10 μg/ml tetracycline to ensure the presence of the F’ episome. A single colony was selected to produce electrocompetent cells with aliquots of the resulting electrocompetent cells used for all experiments within this study. All growth experiments were conducted in LB media supplemented with glucose (2% w/v) and spectinomycin (50 μg/ml) to maintain the pSKunk1 plasmid except where otherwise noted. Expression of the antibiotic resistance proteins was induced by addition of 1 mM IPTG to exponentially growing cultures.

### Library creation

We constructed libraries of all the possible single-codon substitutions in *NDM-1*, *CAT-I*, and *aadB* using inverse PCR with mutagenic oligonucleotides as described in previous work (Mehlhoff and Ostermeier 2020). The oligonucleotides contained a NNN degenerative codon targeted to each codon within the three genes. We constructed the library in three regions for *NDM-1*, three regions for *CAT-I*, and two regions for *aadB* due to read length constraints of Illumina MiSeq. We estimated a minimum of 50,000 transformants would be necessary for each region to have a high probability of having nearly all possible single-codon substitutions (Bosley and Ostermeier 2005). For each region, we repeated the transformation and pooled the resulting colonies until we had an excess of 100,000 transformants. We recovered each library from the LB-agar plates using LB media and glycerol before making aliquots for storage at −80°C.

### Growth competition for measurements of fitness effects

Fitness was measured as described in Mehlhoff *et al*. (Mehlhoff et al. 2020) using a growth competition experiment. Our experiment differed only at the start. Frozen library stocks and wild-type cells were thawed on ice before 50 μl was added to 100 ml media containing 50 μg/ml spectinomycin and 2% w/v glucose in a 500 ml unbaffled shake flask. The cultures were incubated with shaking for 16 hours at 37°C. The wild-type and library cultures were then diluted to an of OD of 0.050 in 100 ml of media and mixed at a ratio of 2:98 based on volume in a fresh unbaffled flask. The cultures were incubated at 37°C with shaking until the OD was about 0.5 and the remainder of the growth competition followed as described in previous work. In short, the exponentially growing library cultures were induced with IPTG and allowed to grow for approximately ten generations with a single dilution at about five generations. Plasmid DNA was collected from before induction and after 10 generations of growth. Custom adapters were added to the plasmid DNA by PCR. Adapters were designed to be compatible with the Illumina platform and contained barcodes for identification of each timepoint and sample. Proper DNA size of the PCR products was validated by agarose electrophoresis gel before the PCR products were pooled and submitted for Illumina MiSeq (2 x 300 bp reads) at the Single Cell and Transcriptomics Core facility at Johns Hopkins University.

### Deep sequencing analysis

Output files from Illumina MiSeq were first run through FastQC (Andrews et al. 2018) to check read quality. The paired-end reads were merged using PEAR (Stamatakis et al. 2014) set to a minimum assembly length of 150 base pairs reads allowing for high quality scores at both ends of the sequence. Adapters were trimmed from the ends of the antibiotic resistance genes’ coding sequence using Trimmomatic (Bolger et al. 2014). Enrich2 (Rubin et al. 2017) was used to count the frequency of each allele for use in calculating selection coefficients and associated statistical measures. We set Enrich2 to filter out any reads containing bases with a quality score below 20, bases marked as N, or mutations at more than one codon.

Fitness of an allele (*w_i_*) was calculated from the enrichment of the synonyms of the wild-type gene (*ε_ωt_*), the enrichment of allele *i* (*ε_*i*_*) and the fold increase in the number of cells during the growth competition experiment (*r*) as described by Equation 1. We utilize the frequency of wildtype synonymous alleles as the reference instead of the frequency of wildtype because wildtype synonyms occurred more frequently in the library and wildtype sequencing counts are more prone to being affected by the artifact of PCR template jumping during the preparation of barcoded amplicons for deep-sequencing. Detailed derivations of the following equations (Equations 1-6) can be found in our previous work (Mehlhoff et al. 2020).

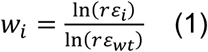

We calculate the variance in the fitness as

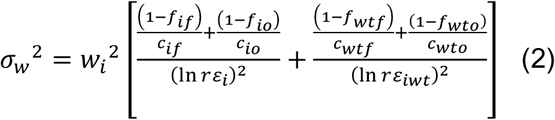

Where the frequency of allele (*f_i_*) is calculated from counts of that allele (*c_i_*) and the total sequencing counts (*c_T_*).

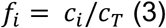

From the variance in fitness, we calculated a 99% confidence interval. Additionally, we calculated a *P*-value using a 2-tailed test. Details of the Z-score and *P*-value equations are available in Mehlhoff *et al*. (Mehlhoff et al. 2020).

We estimated the number of false positives that would be included at *P*<0.01 and *P*<0.001 significance in order to correct for multiple testing (Storey and Tibshirani 2003) in our DMS datasets as described previously (Mehlhoff et al. 2020). For *TEM-1*, we estimated that our data would contain approximately 55.0 false positives on average at *P*<0.01 significance and an estimated 5.6 false positives on average at *P*<0.001 significance for a single replica (Mehlhoff et al. 2020). Those values are 44.1 and 4.3 (*CAT-I*), 52.8 and 5.3 (*NDM-1*), and 33.8 and 3.4 (*aadB*) at *P*<0.01 and *P*<0.001 significance, respectively. We chose to report the frequency of mutations having fitness effects that met the *P-*value criteria in both replica experiments to limit the occurrence of false positives.

### Mutant construction

We constructed a total of 34 mutants across the three genes consisting of 12 *CAT-I* mutants, 13 *NDM-1* mutants, and 9 *aadB* mutants. We used inverse PCR to introduce the mutations. We also used inverse PCR to construct a control plasmid, pSKunk1-ΔGene, which had the coding region of the studied antibiotic resistance genes deleted.

For the C26D and C26S mutants in *NDM-1*, we found that an IS4-like element ISVsa5 family transposase insertion would occur within the *NDM-1* gene during the six hours of induced monoculture growth (Supplementary Text). We made two synonymous mutations within the 5’-GCTGAGC-3’ insertion site that fully overlapped codons 23 and 24 to reduce transposase insertion and get an accurate measure of the collateral fitness effects for the C26D and C26S mutations. The new sequence was 5’-GTTATCA-3’. Inverse PCR was used to introduce these synonymous mutations. All mutant plasmids were transformed into NEB 5-alpha LacI^q^ electrocompetent cells.

### Monoculture growth assay for fitness

Cultures were grown following the same protocol described by Mehlhoff *et al*. (Mehlhoff et al. 2020). Briefly, we tracked the optical density of monocultures over six hours of induced growth with a dilution after three hours. We calculated the fitness using Equation 4 from the dilution factor (*d*) and the starting and final OD’s (*O_o_*, *O_f_*) of the mutant and wild-type cultures under the assumption that the correlation between OD and cell density was the same for cells whether they were expressing wild-type or any mutant TEM-1 allele. Thus, fitness represents the mean growth rate of the cells expressing the mutant protein relative to the growth rate of cells expressing the wildtype protein.

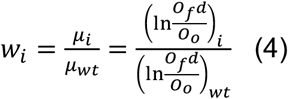

### Cell fractionation and analysis by PAGE and Western blot

Cell cultures were grown as in the monoculture growth assay for fitness and then fractionated following a previously described procedure (Mehlhoff et al. 2020) based on the BugBuster reagent protocol for cell fractionation. Further description of measurement of the resulting protein concentration by DC Assay Kit (Bio-Rad) and sample preparation for SDS-PAGE can be found there as well. We loaded 15 μg of protein from fractionated samples in each lane for SDS-PAGE gels. Protein fractions were analyzed on 12% NuPAGE Bis-Tris gels (Thermo Fisher) in MOPS running buffer. Gels were stained using Coomassie Brilliant Blue R-250 (Bio-Rad) to verify even loading of lanes. We performed immunoblots using rabbit Anti-CAT-I antibody (C9336; Sigma), rabbit Anti-NDM-1 antibody (MBS715336; MyBioSource), and rabbit Anti-AadB antibody (Cusabio custom antibody). All western blot results were substantiated by biological replicates (**supplementary figs. S9, S11**, and **S12**, Supplementary Material online).

Since CAT-I is a homotrimer in its native form (Biswas et al. 2012), we used Native-PAGE and Western blots to see if the quaternary structure of CAT-I was affected by select mutations found to have deleterious collateral fitness effects. Samples for Native-PAGE were prepared using 15 μg of protein, 4X NativePAGE sample buffer (Invitrogen), 0.25 μl NativePAGE 5% G-250 Sample Additive, and nuclease-free water before being loaded onto a NativePAGE 4-16% Bis-Tris Gel (Thermo Fisher). The inner chamber of the electrophoresis system was filled with 1X NativePAGE Dark Blue Cathode Buffer (Thermo Fisher) while the outer chamber was filled with 1X NativePAGE Anode Buffer (Thermo Fisher). The gel was run for approximately two hours at 150 V with the 1X NativePAGE Dark Blue Cathode Buffer being replaced with 1X NativePAGE Light Blue Cathode Buffer (Thermo Fisher) after the dye front had migrated one third of the way through the gel. We then proceeded with Western blots as described.

### Timescale of transposase insertion in C26D and C26S of NDM-1

Monocultures of cells expressing C26D, C26S, and wild-type NDM-1 were grown to an OD of 0.5 before induction following the protocol for monoculture growth assays. We then diluted to an approximate OD of 0.2 and added 1 mM IPTG to each flask. Monocultures were grown for 6 hours with 10 ml of culture collected every hour for plasmid extraction. We diluted cultures back to an OD of 0.2 each hour to maintain exponential growth phase while still having enough cells available at the next hour for plasmid extraction. The resulting DNA was submitted for Sanger sequencing and run on a 1% agarose gel containing ethidium bromide for imaging under UV light.

### Minimum inhibitory concentration (MIC) assays

Cultures were grown overnight at 37°C in LB media containing 2% w/v glucose, 50 μg/ml spectinomycin, and 1 mM IPTG. The overnight cultures were then diluted to an OD of 1×10^−5^ before 50 μl of diluted culture was added to a plate and spread with an L-shaped spreader. Plates were made with 20 ml of LB-agar containing 2% w/v glucose, 50 μg/ml spectinomycin, and 1 mM IPTG. Differing levels of antibiotic were added to each plate depending on the MIC for the wild-type protein. For CAT-I, plates were made with 0, 16, 32, 64, 128, 256, 512, and 1024 μg/ml chloramphenicol. For NDM-1, plates were made with 0, 32, 64, 128, 256, 512, 1024, 2048, and 4096 μg/ml ampicillin. For AadB, plates were made with 0, 8, 16, 32, 64, 128, 256, and 512 μg/ml kanamycin. Plates were incubated for 16 hours at 37°C before colonies were counted. We identified the MIC as the antibiotic concentration at which there were fewer than 5% of the colony forming units (CFU) relative to a plate without antibiotics.

### RNA-Seq Sample Preparation and Analysis

Monocultures expressing the wildtype and all mutants for a given antibiotic resistance gene were grown on the same day along with an uninduced wild-type control. Samples for RNA-seq were prepared as described previously (Mehlhoff et al. 2020). The resulting double-stranded cDNA was pooled and submitted for Illumina TruSeq (2 x 75 bp) at the Single Cell and Transcriptomics Core facility at Johns Hopkins University.

Reads were inspected with FastQC (Andrews et al. 2018) to check per base sequence quality. We indexed the NEB 5-alpha F’Iq genome (NCBI Reference Sequence: NZ_CP053607.1) after having removed genes that appeared in both the episome and chromosomal DNA. Such genes had their counts otherwise overwritten as verified by mapping to the NEB 5-alpha genome (NCBI Reference Sequence: NZ_CP017100.1). We mapped our paired end reads to the indexed NEB 5-alpha F’Iq transcriptome using STAR (Dobin et al. 2013). featureCounts (Liao et al. 2014) was used to quantify and tabulate transcript abundance while edgeR (Robinson et al. 2009) was used to tabulate the normalized counts per million.

### RNA-Seq data analysis

Using Equation 5, we calculated a fold-difference in gene expression (*g*_i_) from the counts of an individual gene from sample *i* (*c*_i_), the counts of that same gene in the corresponding induced wild-type sample (*c*_wt_), and total sequencing counts (*c*_T_).

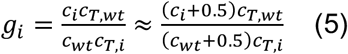

The variance in the fold-difference is then calculated from counts and the fold-difference in gene expression using Equation 6.

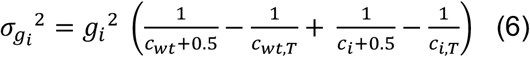

A 99% confidence interval, *Z*-score, and *P*-value were calculated following the method used in our deep sequencing analysis. Full equations are included in Mehlhoff *et al*. (Mehlhoff et al.2020).

We primarily identified genes and more broadly, gene clusters and associated pathways, from the set of values in which there was a >2-fold difference in gene expression with *P*<0.001 significance. We utilized EcoCyc (Keseler et al. 2017) to further identify interconnected genes and search for patterns across genes which potentially play a role in deleterious collateral fitness effects.

### PCR amplification of *traJ* and *traI* genes

We grew monocultures of uninduced and induced wild-type CAT-I along with induced monocultures of the S27M and Q30M mutants following the monoculture growth assay outlined previously. We utilized the Qiagen Large-Construct Kit following the protocol for high yields of large-construct DNA without removal of genomic DNA. The *traJ* and *traI* genes were PCR amplified using 2X GoTaq Green MasterMix starting from 10 ng of template from each sample. PCR products were loaded on a 1% TAE gel containing ethidium bromide. Bands were visualized under UV light and imaged with Carestream Gel Logic 112.

## Acknowledgements

This research was supported by National Science Foundation grants MCB-1817646 and MCB-2113019 to M.O.

## Data availability

DMS sequencing data for *CAT-I*, *NDM-1*, and *aadB* can be found at BioProject (https://www.ncbi.nlm.nih.gov/bioproject) using accession no. PRJNA818255. RNA-Seq data are available at Gene Expression Omnibus at NCBI (https://www.ncbi.nlm.nih.gov/geo/) using accession no. GSE199248. Sequencing counts for the DMS studies can be found in **supplementary data S1-3**, Supplementary Material online. Sequencing counts for the RNA-Seq experiment can be found in **supplementary data S4**, Supplementary Material online.

## References

Agozzino L, Dill KA. 2018. Protein evolution speed depends on its stability and abundance and on chaperone concentrations. Proc. Natl. Acad. Sci. U. S. A. 115:9092–9097.

Alvarez-Ponce D, Aguilar-Rodríguez J, Fares MA, Papp B. 2019. Molecular Chaperones Accelerate the Evolution of Their Protein Clients in Yeast. Genome Biol. Evol. 11:2360–2375.

Andrews S, Wingett SW, Hamilton RS. 2018. FastQ Screen: A tool for multi-genome mapping and quality control F1000Res 7:1338.

Auclair SM, Bhanu MK, Kendall DA. 2012. Signal peptidase I: Cleaving the way to mature proteins. Protein Sci. 21:13–25.

Awad A, Arafat N, Elhadidy M. 2016. Genetic elements associated with antimicrobial resistance among avian pathogenic Escherichia coli. Ann. Clin. Microbiol. Antimicrob. 15:1–8.

Bahr G, Vitor-Horen L, Bethel CR, Bonomo RA, Gonzalez LJ, Vila AJ. 2018. Clinical Evolution of New Delhi Metallo-β-Lactamase (NDM) optimizes resistance under Zn(II) Deprivation. Antimicrob. Agents Chemother. 62:e01849–17.

De Biase D, Tramonti A, Bossa F, Visca P. 1999. The response to stationary-phase stress conditions in Escherichia coli: role and regulation of the glutamic acid decarboxylase system. Mol. Microbiol. 32:1198–1211.

Biesiadecka MK, Sliwa P, Tomala K, Korona R. 2020. An Overexpression Experiment Does Not Support the Hypothesis That Avoidance of Toxicity Determines the Rate of Protein Evolution. Genome Biol. Evol. 12:589–596.

Biswas T, Houghton JL, Garneau-Tsodikova S, Tsodikov O V. 2012. The structural basis for substrate versatility of chloramphenicol acetyltransferase CAT I. Protein Sci. 21:520–530.

Bolger AM, Lohse M, Usadel B. 2014. Trimmomatic: a flexible trimmer for Illumina sequence data. Bioinformatics 30:2114–2120.

Bosley AD, Ostermeier M. 2005. Mathematical expressions useful in the construction, description and evaluation of protein libraries. Biomol. Eng. 22:57–61.

Bratulic S, Gerber F, Wagner A. 2015. Mistranslation drives the evolution of robustness in TEM-1 β-lactamase. Proc. Natl. Acad. Sci. U. S. A. 112:12758–12763.

Cheng Z, Thomas PW, Ju L, Bergstrom A, Mason K, Clayton D, Miller C, Bethel CR, VanPelt J, Tierney DL, et al. 2018. Evolution of New Delhi metallo-β-lactamase (NDM) in the clinic: Effects of NDM mutations on stability, zinc affinity, and mono-zinc activity. J. Biol. Chem. 293:12606–12618.

Cox G, Stogios PJ, Savchenko A, Wright GD. 2015. Structural and molecular basis for resistance to aminoglycoside antibiotics by the adenylyltransferase ANT(2″)-Ia. MBio 6:1–9.

Dobin A, Davis CA, Schlesinger F, Drenkow J, Zaleski C, Jha S, Batut P, Chaisson M, Gingeras TR. 2013. STAR - ultrafast universal RNA-seq aligner. Bioinformatics 29:15–21.

Firnberg E, Ostermeier M. 2012. PFunkel: Efficient, Expansive, User-Defined Mutagenesis. PLoS One 7:1–10.

Flores-Kim J, Darwin AJ. 2016. The phage shock protein response. Annu. Rev. Microbiol. 70:83–101.

Geiler-Samerotte KA, Dion MF, Budnik BA, Wang SM, Hartl DL, Drummond DA. 2011. Misfolded proteins impose a dosage-dependent fitness cost and trigger a cytosolic unfolded protein response in yeast. Proc. Natl. Acad. Sci. U. S. A. 108:680–685.

González LJ, Bahr G, Nakashige TG, Nolan EM, Bonomo RA, Vila AJ. 2016. Membrane anchoring stabilizes and favors secretion of New Delhi metallo-β-lactamase. Nat. Chem. Biol. 12:516–522.

Goodale A, Michailidis F, Watts R, Chok SC, Hayes F. 2020. Characterization of permissive and non-permissive peptide insertion sites in chloramphenicol acetyltransferase. Microb. Pathog. 149:104395.

Grabowicz M, Silhavy TJ. 2017. Envelope stress responses: an interconnected safety net. Trends Biochem. Sci. 42:232–242.

Guo Y, Wang J, Niu G, Shui W, Sun Y, Zhou H, Zhang Y, Yang C, Lou Z, Rao Z. 2011. A structural view of the antibiotic degradation enzyme NDM-1 from a superbug. Protein Cell 2:384–394.

Halling SM, Kleckner N. 1982. A symmetrical six-base-pair target site sequence determines Tn10 insertion specificity. Cell 28:155–163.

Holden ER, Webber MA. 2020. MarA, RamA, and SoxS as mediators of the stress response: survival at a cost. Front. Microbiol. 11:1–10.

Jahn TR, Radford SE. 2008. Folding versus aggregation: polypeptide conformations on competing pathways. Arch. Biochem. Biophys. 469:100–117.

Kaminska R, Van Der Woude MW. 2010. Establishing and maintaining sequestration of dam target sites for phase variation of agn43 in Escherichia coli. J. Bacteriol. 192:1937–1945.

Keseler IM, Mackie A, Santos-Zavaleta A, Billington R, Bonavides-Martínez C, Caspi R, Fulcher C, Gama-Castro S, Kothari A, Krummenacker M, et al. 2017. The EcoCyc database: reflecting new knowledge about Escherichia coli K-12. Nucleic Acids Res. 45:D543–D550.

King D, Strynadka N. 2011. Crystal structure of New Delhi metallo-β-lactamase reveals molecular basis for antibiotic resistance. Protein Sci. 20:1484–1491.

Krylov DM, Wolf YI, Rogozin IB, Koonin E V. 2003. Gene loss, protein sequence divergence, gene dispensability, expression level, and interactivity are correlated in eukaryotic evolution. Genome Res. 13:2229–2235.

Laganenka L, Colin R, Sourjik V. 2016. Chemotaxis towards autoinducer 2 mediates autoaggregation in Escherichia coli. Nat. Commun. 7:1–10.

Lemos B, Bettencourt BR, Meiklejohn CD, Hartl DL. 2005. Evolution of proteins and gene expression levels are coupled in Drosophila and are independently associated with mRNA abundance, protein length, and number of protein-protein interactions. Mol. Biol. Evol. 22:1345–1354.

Levy ED, De S, Teichmann SA. 2012. Cellular crowding imposes global constraints on the chemistry and evolution of proteomes. Proc. Natl. Acad. Sci. U. S. A. 109:20461–20466.

Liao Y, Smyth GK, Shi W. 2014. FeatureCounts: An efficient general purpose program for assigning sequence reads to genomic features. Bioinformatics 30:923–930.

Mehlhoff JD, Ostermeier M. 2020. Biological fitness landscapes by deep mutational scanning. Methods Enzymol. 643:203–224.

Mehlhoff JD, Stearns FW, Rohm D, Wang B, Tsou E, Dutta N, Hsiao M-H, Gonzalez CE, Rubin AF, Ostermeier M. 2020. Collateral fitness effects of mutations. Proc. Natl. Acad. Sci. 117:11597–11607.

Mironov A, Seregina T, Shatalin K, Nagornykh M, Shakulov R, Nudler E. 2020. CydDC functions as a cytoplasmic cystine reductase to sensitize Escherichia coli to oxidative stress and aminoglycosides. Proc. Natl. Acad. Sci. U. S. A. 117:23565–53570.

Navarro S, Villar-Piqué A, Ventura S. 2014. Selection against toxic aggregation-prone protein sequences in bacteria. Biochim. Biophys. Acta - Mol. Cell Res. 1843:866–874.

Olzscha H, Schermann SM, Woerner AC, Pinkert S, Hecht MH, Tartaglia GG, Vendruscolo M, Hayer-Hartl M, Hartl FU, Vabulas RM. 2011. Amyloid-like aggregates sequester numerous metastable proteins with essential cellular functions. Cell 144:67–78.

Parsell DA, Sauer RT. 1989. Induction of a heat shock-like response by unfolded protein in Escherichia coli: dependence on protein level not protein degradation. Genes Dev. 3:1226–1232.

Peralta DR, Adler C, Corbalán NS, Paz García EC, Pomares MF, Vincent PA. 2016. Enterobactin as part of the oxidative stress response repertoire. PLoS One 11:1–15.

Prunotto A, Bahr G, González LJ, Vila AJ, Peraro MD. 2020. Molecular bases of the membrane association mechanism potentiating antibiotic resistance by New Delhi metallo-β-Lactamase 1. ACS Infect. Dis. 6:2719–2731.

Raivio TL. 2014. Everything old is new again: An update on current research on the Cpx envelope stress response. Biochim. Biophys. Acta - Mol. Cell Res. 1843:1529–1541.

Reis AC, Salis HM. 2020. An automated model test system for systematic development and improvement of gene expression models. ACS Synth. Biol. 9:3145–3156.

Robinson MD, McCarthy DJ, Smyth GK. 2009. edgeR: A Bioconductor package for differential expression analysis of digital gene expression data. Bioinformatics 26:139–140.

Roth M, Jaquet V, Lemeille S, Bonetti E-J, Cambet Y, François P, Krause K-H. 2022. Transcriptomic analysis of E. coli after exposure to a sublethal concentration of hydrogen peroxide revealed a coordinated up-regulation of the cysteine biosynthesis pathway. Antioxidants (Basel) 11:655.

Rubin AF, Gelman H, Lucas N, Bajjalieh SM, Papenfuss AT, Speed TP, Fowler DM. 2017. A statistical framework for analyzing deep mutational scanning data. Genome Biol. 19:17.

Van der Schueren J, Robben J, Goossens K, Heremans K, Volckaert G. 1996. Identification of local carboxy-terminal hydrophobic interactions essential for folding or stability of chloramphenicol acetyltransferase. J. Mol. Biol. 256:878–888.

Sohka T, Heins RA, Phelan RM, Greisler JM, Townsend CA, Ostermeier M. 2009. An externally tunable bacterial band-pass filter. Proc. Natl. Acad. Sci. 106:10135–10140.

Stamatakis A, Zhang J, Kobert K. 2014. PEAR: a fast and accurate Illumina Paired-End reAd mergeR. Bioinformatics 30:614–620.

Storey JD, Tibshirani R. 2003. Statistical significance for genomewide studies. Proc. Natl. Acad. Sci. U. S. A. 100:9440–5.

Sun Z, Hu L, Sankaran B, Prasad BVV, Palzkill T. 2018. Differential active site requirements for NDM-1 β-lactamase hydrolysis of carbapenem versus penicillin and cephalosporin antibiotics. Nat. Commun. 9:4524.

Taverna DM, Goldstein RA. 2002. Why are proteins marginally stable? Proteins 46:105–109.

Thomas PW, Zheng M, Wu S, Guo H, Liu D, Xu D, Fast W. 2011. Characterization of purified New Delhi metallo-β-lactamase-1. Biochemistry 50:10102–10113.

Tucker DL, Foster JW, Miranda RL, Cohen PS, Conway T. 2003. Genes of the GadX-GadW regulon J. Bacteriol. 185:3190–201.

Usmanova DR, Plata G, Vitkup D. 2021. The relationship between the misfolding avoidance hypothesis and protein evolutionary rates in the light of empirical evidence. Genome Biol. Evol. 13:evab006.

Wall E, Majdalani N, Gottesman S. 2018. The complex Rcs regulatory cascade. Annu. Rev. Microbiol. 72:111–139.

Wright E, Serpersu EH. 2005. Enzyme-substrate interactions with an antibiotic resistance enzyme: Aminoglycoside nucleotidyltransferase(2″)-Ia characterized by kinetic and thermodynamic methods. Biochemistry 44:11581–11591.

Wu Z, Cai X, Zhang X, Liu Y, Tian G-B, Yang J-R, Chen X. 2022. Expression level is a major modifier of the fitness landscape of a protein coding gene. Nat. Ecol. Evol. 6:103–115.

Yang JR, Liao BY, Zhuang SM, Zhang J. 2012. Protein misinteraction avoidance causes highly expressed proteins to evolve slowly. Proc. Natl. Acad. Sci. U. S. A. 109:831–840.

Yong D, Toleman MA, Giske CG, Cho HS, Sundman K, Lee K, Walsh TR. 2009. Characterization of a new metallo-β-lactamase gene, bla NDM-1, and a novel erythromycin esterase gene carried on a unique genetic structure in Klebsiella pneumoniae sequence type 14 from India. Antimicrob. Agents Chemother. 53:5046–5054.

Zalucki YM, Jen FEC, Pegg CL, Nouwens AS, Schulz BL, Jennings MP. 2020. Evolution for improved secretion and fitness may be the selective pressures leading to the emergence of two NDM alleles. Biochem. Biophys. Res. Commun. 524:555–560.

Zhang J, Yang JR. 2015. Determinants of the rate of protein sequence evolution. Nat. Rev. Genet. 16:409–420.

