## Supplemental Text, Figures, and Tables for "Genes vary greatly in their propensity for collateral fitness effects of mutations"

**This PDF file includes:**

Supplementary Text
Figs. S1 to S17
Table S1 to S2
Supplementary References
Captions for Data S1 to S5

**Other Supplementary Materials for this manuscript include the following:**

Data S1 to S5

### Supplementary Text

#### Artifactual fitness values at select residues in NDM-1 and AadB

Although we do not rule out the possibility of beneficial collateral fitness effects in some circumstances, we think they are unlikely to be of sufficient magnitude for us to observed them in our genes under the examined expression levels. From the DMS studies for *NDM-1*, we measured apparent beneficial fitness effects in both biological replicates at residues 26-28 (**supplementary fig. S3A-C**, Supplementary Material online). For residues 132-134 we observed widespread deleterious fitness effects in the first replicate, but widespread beneficial effects of approximately 4% in replica 2. For *aadB*, we measured widespread beneficial fitness effects of approximately 3-6% at residues 155 and 156 for both replicates of DMS (**supplementary fig. S4A,B**, Supplementary Material online). We believe these systematic beneficial fitness effects at these positions are artifactual and not true beneficial collateral fitness effects. Foremost in our reasoning, these beneficial fitness effects in *NDM-1* and *aadB* were not reproducible by monoculture growth experiments for the selected mutants that we tested (**supplementary fig. S5**, Supplementary Material online). Furthermore, these apparent fitness effects are near-universal in the DMS measurements across all substitutions at these positions including nonsense mutations. If nonsense mutations were in fact alleviating the effects of expressing a toxic protein, we would expect to see more widespread beneficial effects at nonsense mutations throughout the protein or beneficial effects from mutations at the start codon. We do not find evidence of either in the DMS data or growth studies with cells expressing *NDM-1* and *aadB*. In addition, the sequencing counts at these positions tend to have abnormally high counts and/or result in abnormally high standard deviations relative to the mean. This can be seen by plotting the ratio between the standard deviation of fitness effects at a position and the mean fitness at that position as a function of total counts. We expect the ratio to increase due to noise at low total count values. At high values of total counts, the ratio should go to a limiting, finite value. We find residues 26-28 and 132-134 in *NDM-1* to be outliers in this plot (**supplementary fig. S3D**, Supplementary Material online). Similarly, residue 156 stands out as an outlier in the plot for *aadB* (**supplementary fig. S4C**, Supplementary Material online). We speculate these artifacts stem from sequencing artifacts and/or with the creation of the amplicons by PCR, but we do not have a specific hypothesis. Because of our lack of confidence in these effects, we do not include them in our analysis. We removed the wild-type synonym counts for residues A155 and C156 of *aadB* region 2 and C26 and P28 of *NDM-1* region 1 so that they would not affect the fitness values calculated when using the wild-type synonym alleles as a reference.

#### Additional changes in gene expression.

We enumerate additional genes we identified whose expression was affected by mutations causing collateral fitness effects.

Expression of genes corresponding to biofilm formation increased for many of the deleterious NDM-1 mutations while genes corresponding to pilus formation, notably belonging to the *fim* and *ecp* operons, decreased in expression (**supplementary fig. S16A**, Supplementary Material online). We note the increase in expression for many of these genes alongside those belonging to the Rcs pathway as further evidence of the biofilm response seen in deleterious mutants.

In contrast, many of the cold-shock proteins are expressed at reduced levels in cells expressing deleterious mutations (**supplementary fig. S16B**, Supplementary Material online). Absence of proteins coded by the *cspIBF* operon results in a competitive disadvantage in stationary phase (Kram et al. 2020). We find that the cultures with higher ODs at the end of the growth period tend to have increased expression of these cold-shock proteins and believe they may be beginning to transition to stationary phase.

We also find increased expression of genes belonging to the fructose catabolic process within cells expressing deleterious mutations of *NDM-1*, *CAT-I*, and *aadB* (**supplementary fig. S16C**, Supplementary Material online) although we do not have a proposed mechanism by which these mutations would lead to such increase in expression.

In our previous RNA-Seq experiments with TEM-1, we noted some samples displayed differential expression for genes regulated by FNR. The FNR-regulated operons include genes which are involved in anaerobic respiration (Partridge et al. 2007) along with the transition from aerobic to micro-anaerobic growth conditions in the presence of nitrate (Constantinidou et al. 2006). We find some of these same differential expression patterns in our RNA-seq experiments for the current study (**supplementary fig. S17**, Supplementary Material online). However, such effects are mainly limited to select samples from the CAT-I gene. We suspect the variability across samples pertains to the availability of oxygen during the processing steps between the end of the monoculture growth assay and RNA extraction. We did not find the differential expression for FNR-regulated operons to impact gene expression in the stress-response pathways.

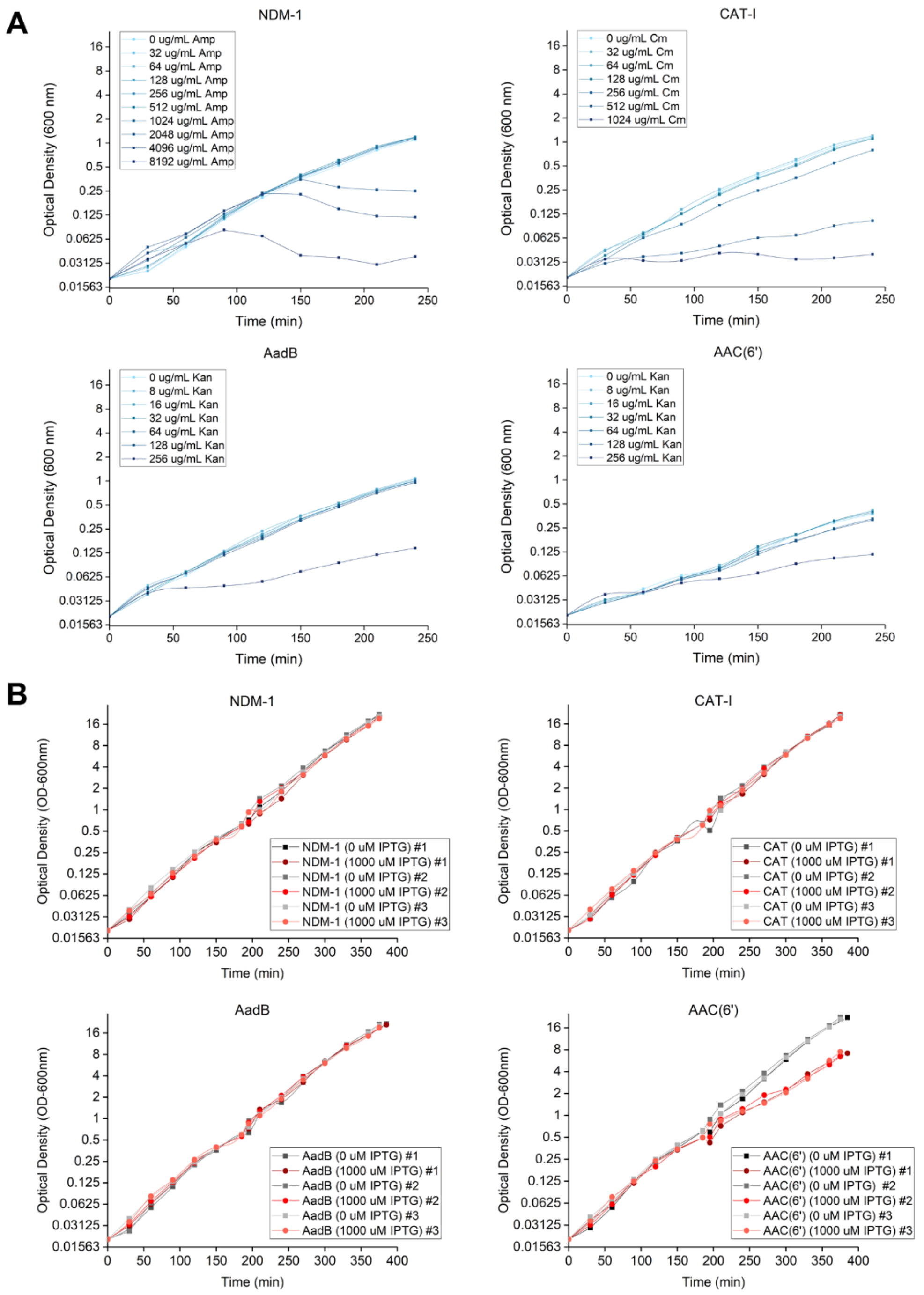


**Figure S1. Effect of expressing antibiotic resistance genes.** (**A**) Confirmation of conferral of antibiotic resistance. Antibiotics were added at the indicated concentrations to exponentially growing cells expressing the *NDM-1*, *CAT*-I, *aadB*, or *aac(6')-Im* antibiotic resistance genes in order to verify their ability to confer antibiotic resistance and assess the cells level of resistance. (**B**) Fitness effect of expression of the antibiotic resistance genes. At time zero, exponentially growing cultures were diluted in two shake flasks to an OD of 0.020 and final volume of 100 ml using LB media which had been prewarmed to 37°C. IPTG was introduced to one of the flasks at a concentration of 1 mM. No IPTG was added to the second flask. Both flasks were incubated at 37°C with shaking and optical density at 600 nm (OD-600 nm) was monitored every 30 minutes for six hours. After approximately three hours, the culture was diluted into fresh LB media (with or without IPTG as appropriate) to maintain growth in exponential phase. The OD measurements after this dilution are relative values in which the measured OD value is multiplied by the dilution factor. From the OD at the start and end of growth and the number of generations, we calculated a fitness effect for expressing each of the antibiotic resistance genes. The average fitness effects of expression in three replica experiments were +1.06% ± 0.06%, -0.33 ± 0.56%, -0.95 ± 0.33%, and -16.14 ± 1.15% for *NDM-1*, *CAT-I*, *aadB*, and *aac(6')-Im*, respectively.

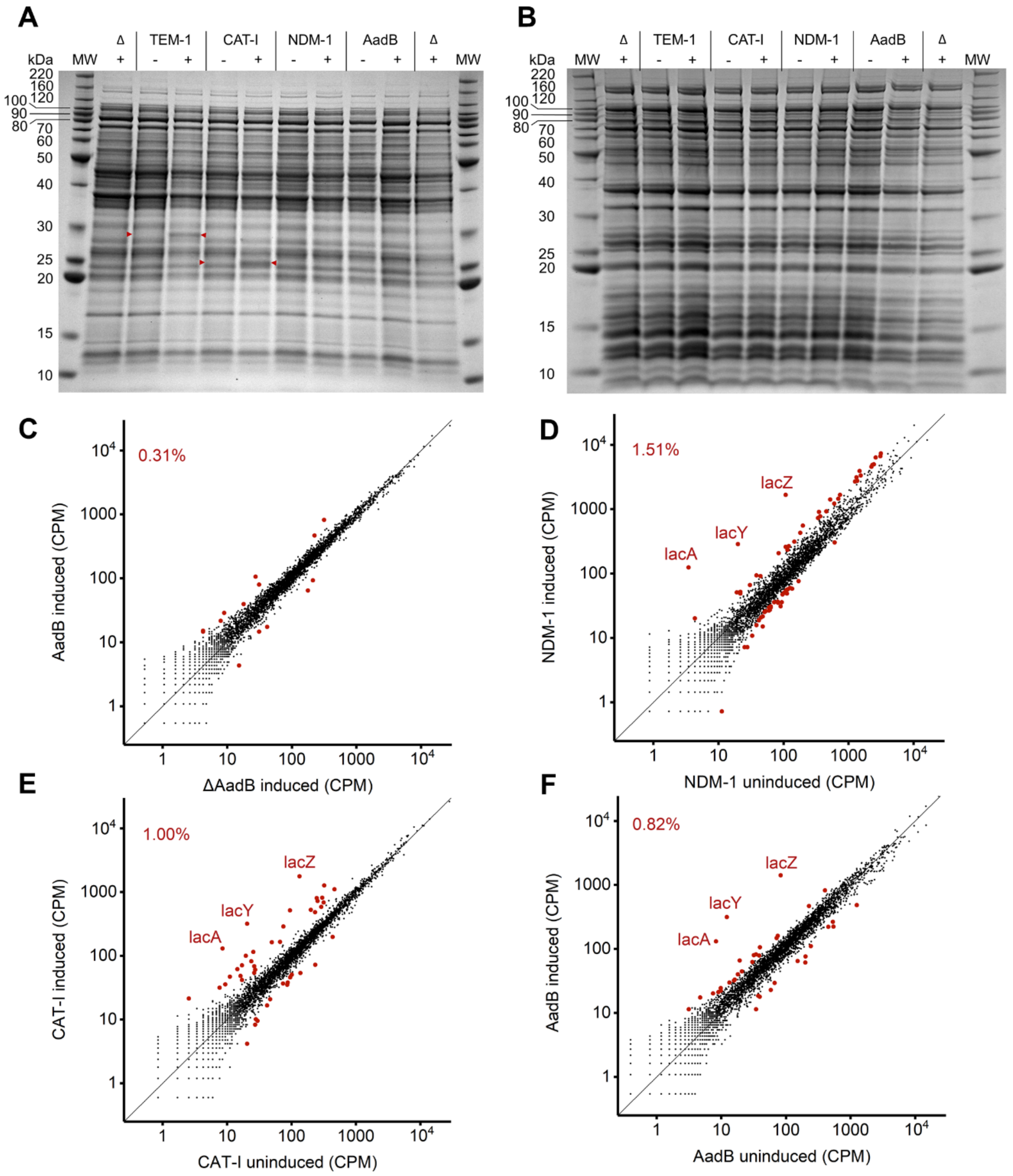


**Figure S2. Expression of wild-type antibiotic resistance proteins and their effects.** SDS-PAGE gel of cell extracts of (**A**) total soluble protein and (**B**) total insoluble protein from NEB 5-alpha F’lacI$^q$ *E. coli* cells containing pSKunk-1-TEM-1, pSKunk-1-CAT-I, pSKunk-1-NDM-1, pSKunk-1-AadB, or the Δ gene control which has the antibiotic resistance gene deleted from the plasmid. Expression of the antibiotic resistance genes was controlled under the *tac* promoter and either uninduced (-) or induced (+) with 1 mM IPTG. Visible changes in expression of a protein band at the expected size of the expressed protein are marked with a red triangle. Bands for NDM-1 and AadB were not observed, but western blots confirmed expression. The expected sizes of the antibiotic resistance proteins are 28.9 kDa for mature TEM-1, 25.7 kDa for CAT-I, 25.9 kDa for mature NDM-1, and 19.9 kDa for AadB. (**C**) RNA-Seq results showing the effect of AadB expression on the *E. coli* transcriptome using Δ gene plasmid as the comparison. RNA-Seq results comparing gene expression in *E. coli* cells expressing (**D**) NDM-1, (**E**) CAT-I, and (**F**) AadB relative to cultures lacking the IPTG inducer. IPTG induces expression of the *lacZYA* operon. Genes with at least a 2-fold change in expression at *P*<0.001 (*Z*-test) significance are indicated with a larger, red circle and the percentage of genes that meet these criteria is indicated on each plot. CPM = counts per million.

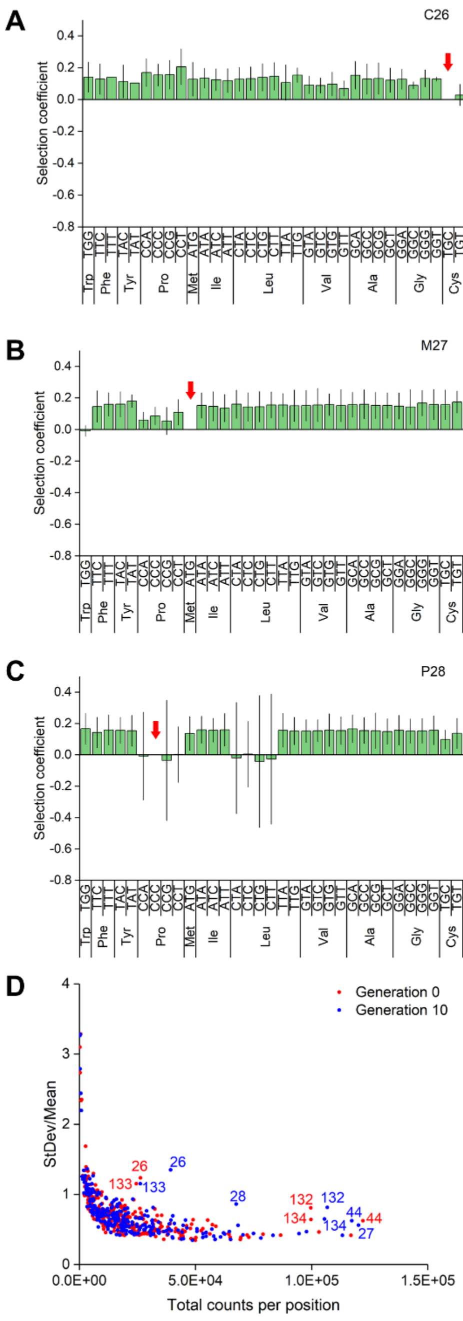


**Figure S3. Artifactual beneficial fitness effects at residues 26-28 of NDM-1.** Weighted mean collateral fitness effects of mutations in (**A**) C26, (**B**) M27, and (**C**) P28 as measured by two replica DMS growth competition experiments. Red arrow indicates wild-type codon. Error bars are 99% confidence intervals. (**D**) Plot of the standard deviation of all fitness measurements for nonsynonymous mutations at a specific residue divided by the mean fitness at that residue as a function of the total counts at the position. Data in red represents counts of a specific allele from before induction and data in blue represents counts of an allele after ten generations of growth.

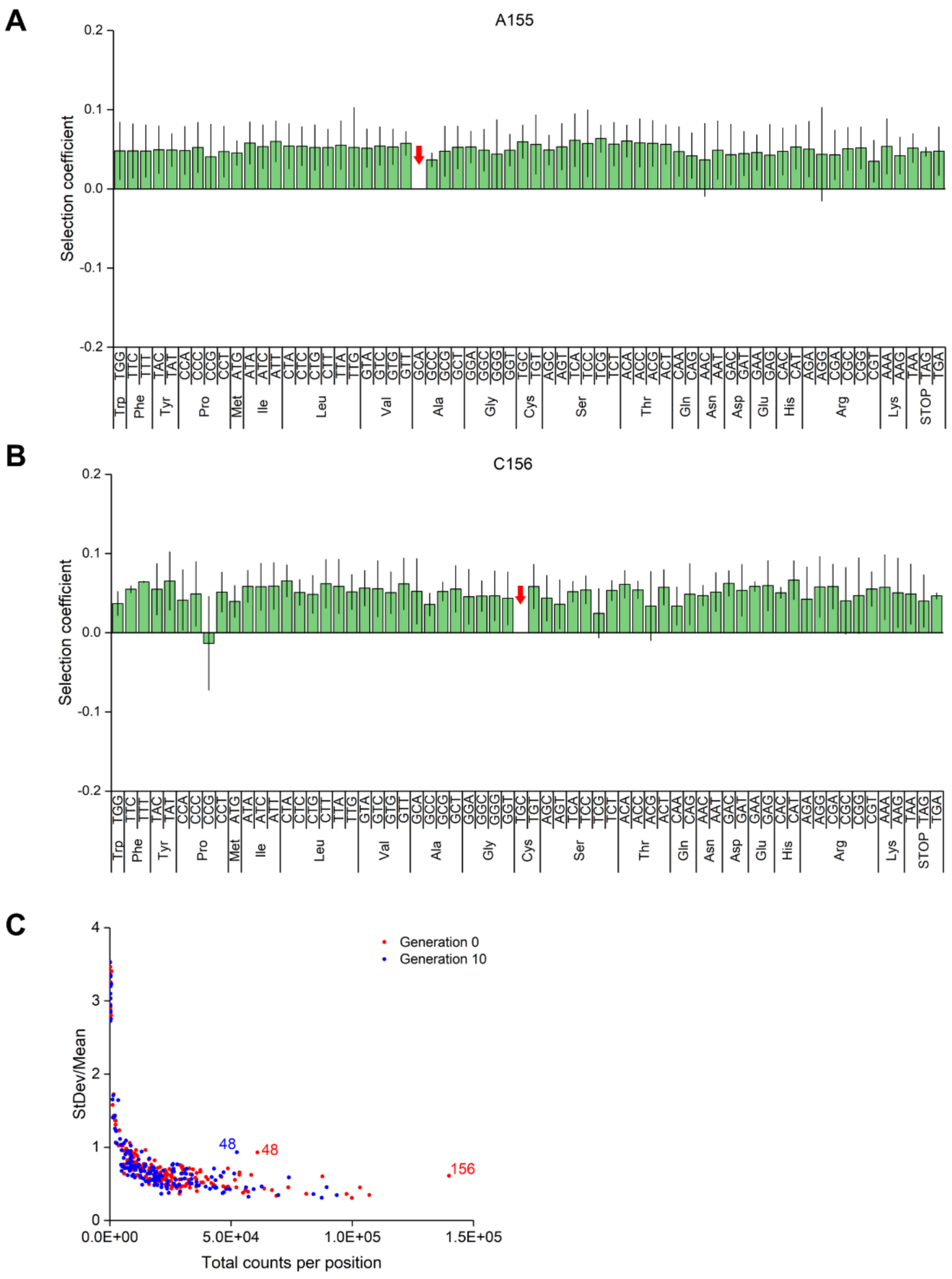


**Figure S4. Artifactual beneficial fitness effects at residues 155 and 156 of AadB.** Weighted mean collateral fitness effects of mutations in (**A**) A155 and (**B**) C156 as measured by two replica DMS growth competition experiments. Red arrow indicates wild-type codon. Error bars are 99% confidence intervals. (**C**) Plot of the standard deviation of all fitness measurements for nonsynonymous mutations at a specific residue divided by the mean fitness at that residue as a function of the total counts at the position. Data in red represents counts of a specific allele from before induction and data in blue represents counts of an allele after ten generations of growth.

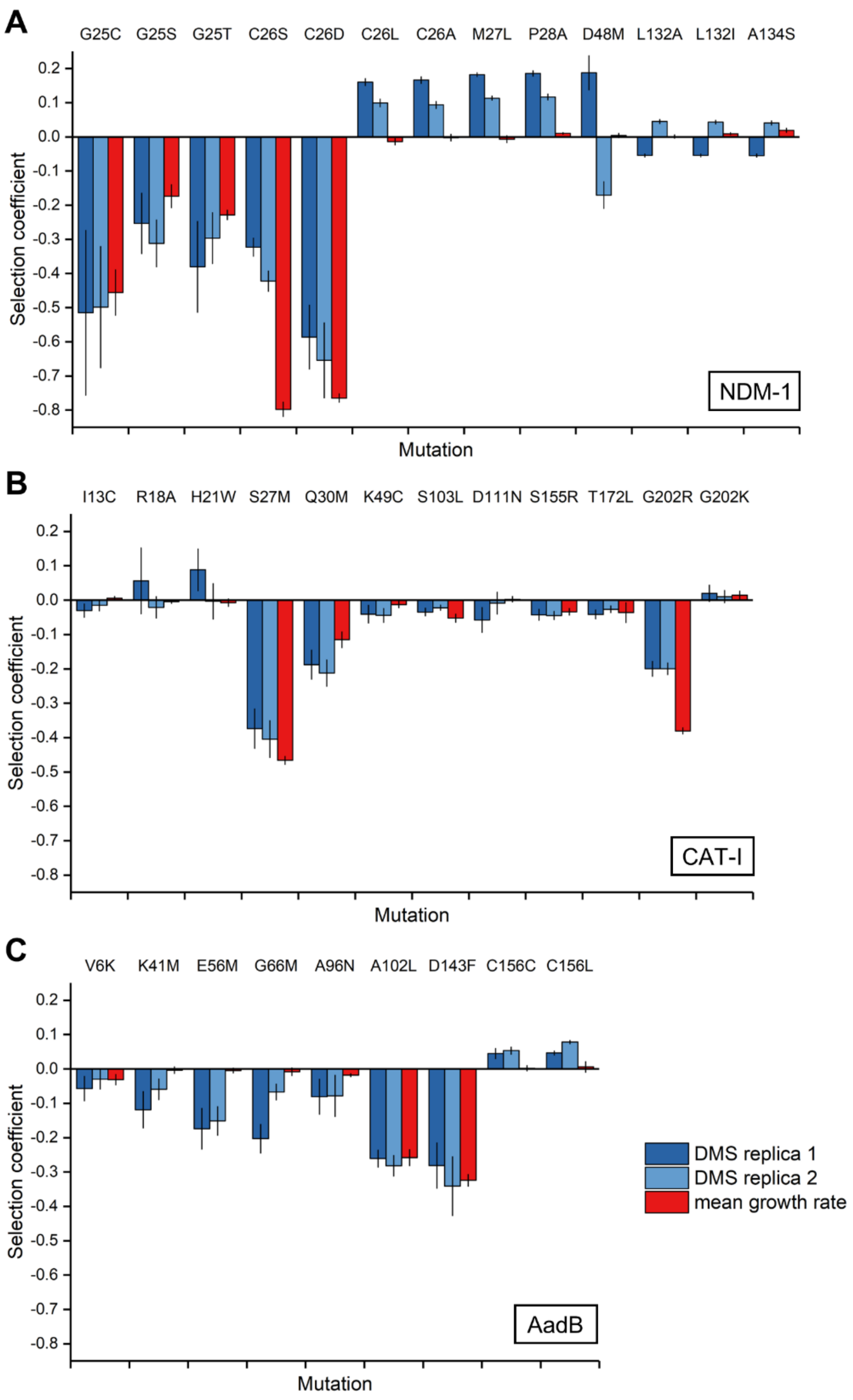


**Figure S5. Confirmation of collateral fitness effects via monoculture growth.** Selection coefficients for cells expressing mutations in (**A**) NDM-1, (**B**) CAT-**I**, and (**C**) AadB as measured by DMS (blue, two biological replicates) and mean growth rate over 6 h postinduction (red; *n* = 3). Mean growth rate was calculated from the starting and final OD (600 nm) for cells expressing wild-type and mutant versions of the antibiotic resistance proteins. Cells were grown in monoculture for 10 generations of wild-type growth. Error bars represent 99% confidence intervals. The C26S and C26D mutants of *NDM-1* contain synonymous mutations at L23 and S24 to eliminate the transposase insertion site. The C156C mutation in *aadB* is a synonymous mutation from TGC to TGT.

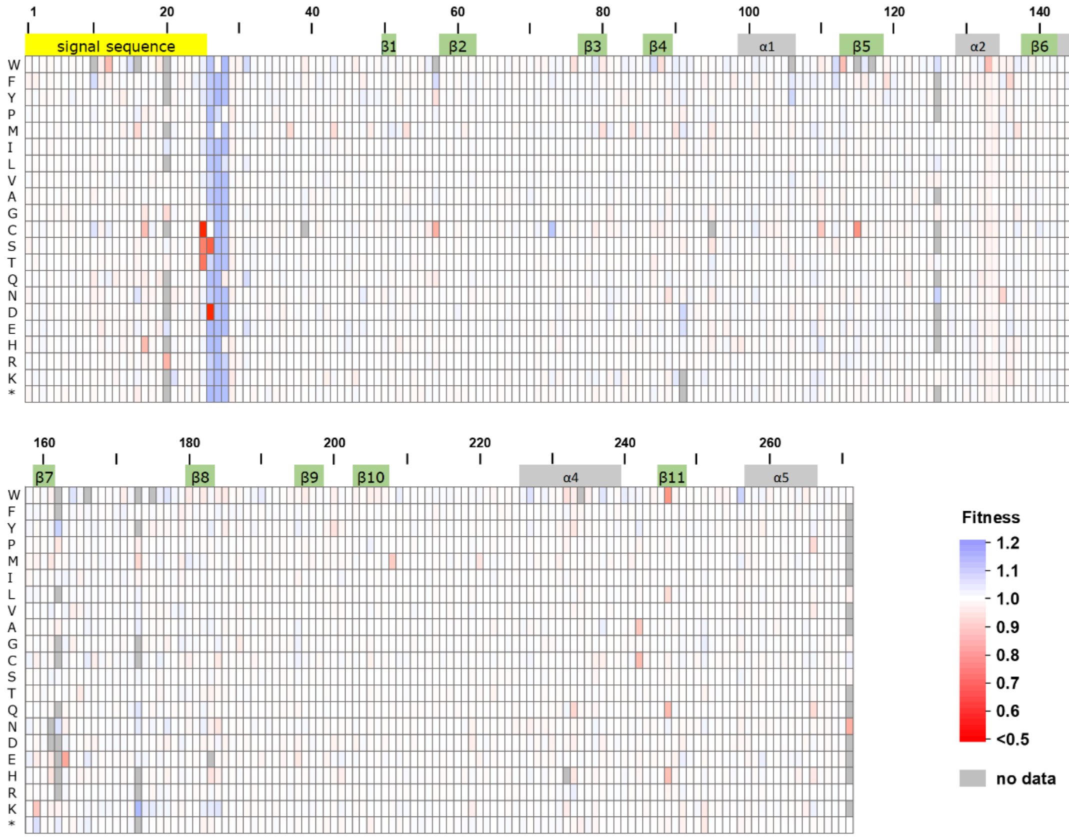


**Figure S6. Weighted mean fitness measurements for NDM-1.** Heat map of the weighted mean collateral fitness effects from two biological replicates of DMS in NDM-1. Cells expressing NDM-1 with the indicated missense or nonsense mutations were grown at 37°C for approximately 10 generations in LB media supplemented with 2.0% glucose and 1 mM IPTG. Mutations missing a fitness measurement in one or both replicates are shaded grey for no data. Regions corresponding to the signal sequence (yellow), α-helices (grey), and β-strands (green) are labeled along the top of the landscape. Tabulated fitness values are provided as **supplementary data S1**, Supplementary Material online.

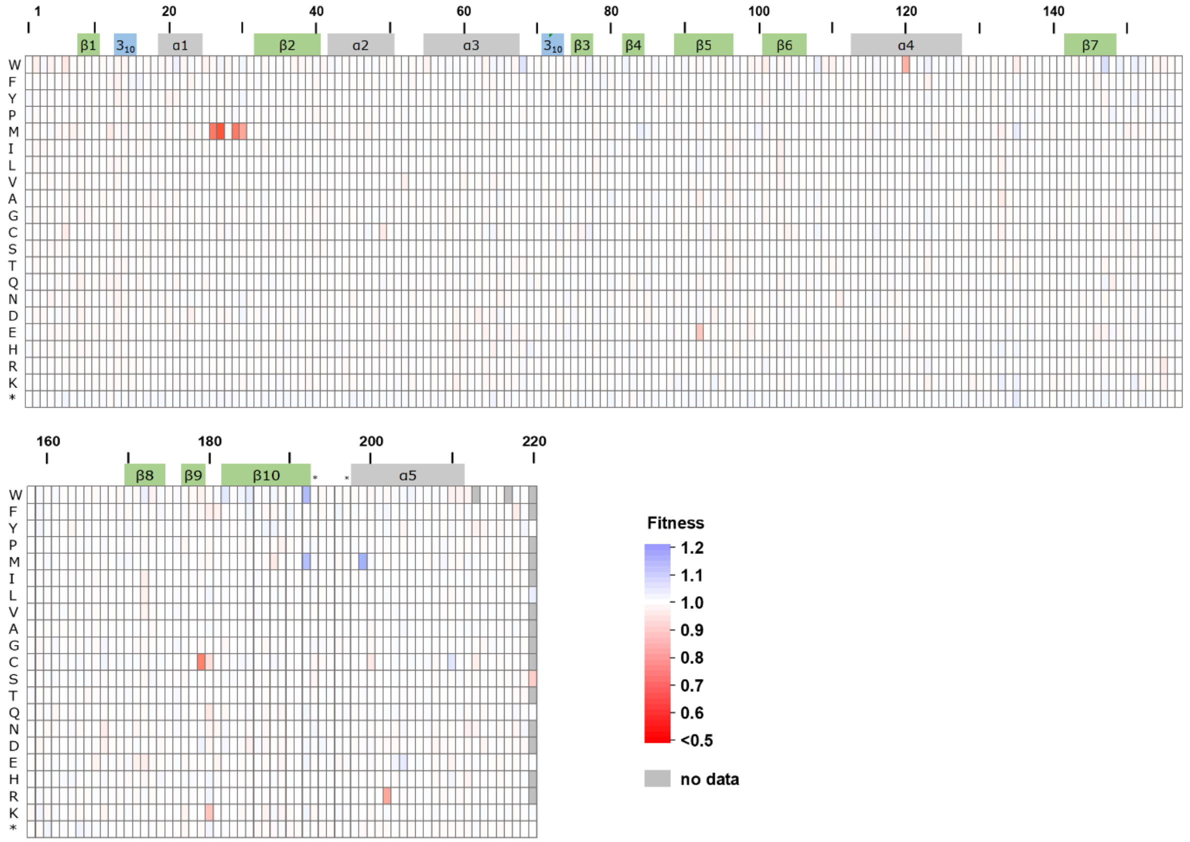


**Figure S7. Weighted mean fitness measurements for CAT-I.** Heat map of the weighted mean collateral fitness effects from two biological replicates of DMS in CAT-I. Cells expressing CAT-I with the indicated missense or nonsense mutations were grown at 37°C for approximately 10 generations in LB media supplemented with 2.0% glucose and 1 mM IPTG. Mutations missing a fitness measurement in one or both replicates are marked in grey. Regions corresponding to α-helices (grey), β-strands (green), $3_{10}$-helices (blue), and key active site residues (*) are labeled along the top of the landscape. Tabulated fitness values are provided as **supplementary data S2**, Supplementary Material online.

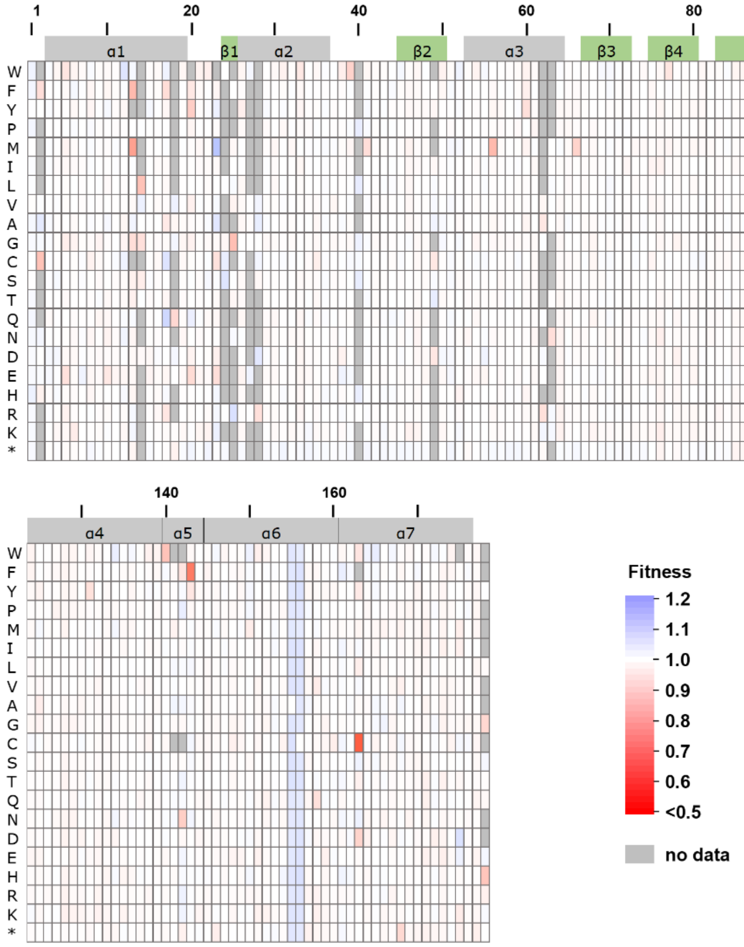


**Figure S8. Weighted mean fitness measurements for AadB.** Heat map of the weighted mean collateral fitness effects from two biological replicates of DMS in AadB. Cells expressing AadB with the indicated missense or nonsense mutations were grown at 37°C for approximately 10 generations in LB media supplemented with 2.0% glucose and 1 mM IPTG. Mutations missing a fitness measurement in one or both replicates are marked in grey. Regions corresponding to α-helices (grey) and β-strands (green) are labeled along the top of the landscape. Tabulated fitness values are provided as **supplementary data S3**, Supplementary Material online.

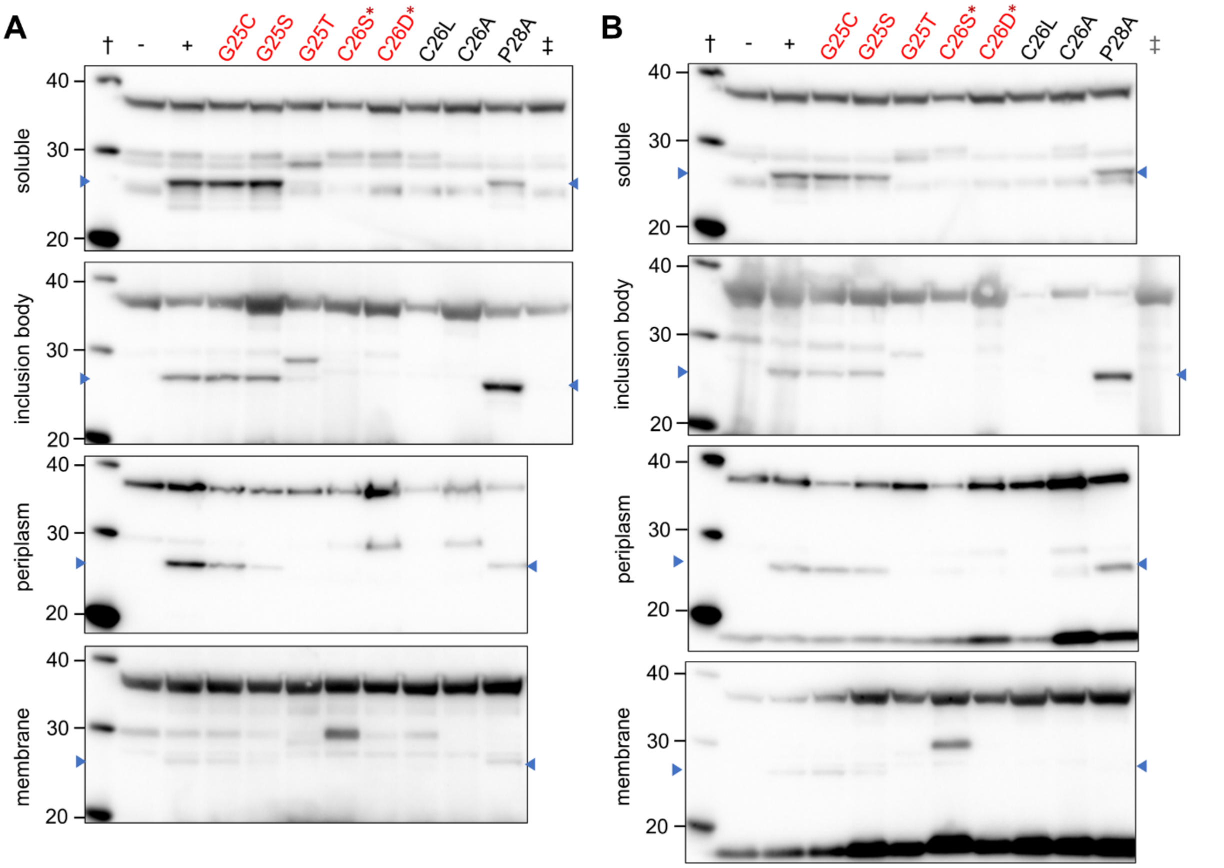


**Figure S9. Western blots of fractionated cells expressing mutants of NDM-1.** Cells expressing NDM-1 over six hours of induced growth were harvested, lysed, and fractionated into total soluble, inclusion body, periplasmic, and membrane fractions. Western blots for the (**A**) first and (**B**) second biological replicate are shown with deleterious mutations labeled in red and neutral mutations labeled in black. Samples from fractionated cells without induction of wild-type NDM-1 (-) and cells expressing wild-type NDM-1 (+) are included on the left of each blot along with the molecular weight standard (†). The C26S and C26D mutants include two synonymous mutations to prevent a transposase insertion indicated by (*). *CAT-I* samples (‡) corresponding to the respective fraction were included on Western blots to identify bands for *E. coli* proteins cross-reacting with the anti-NDM-1 antibodies.

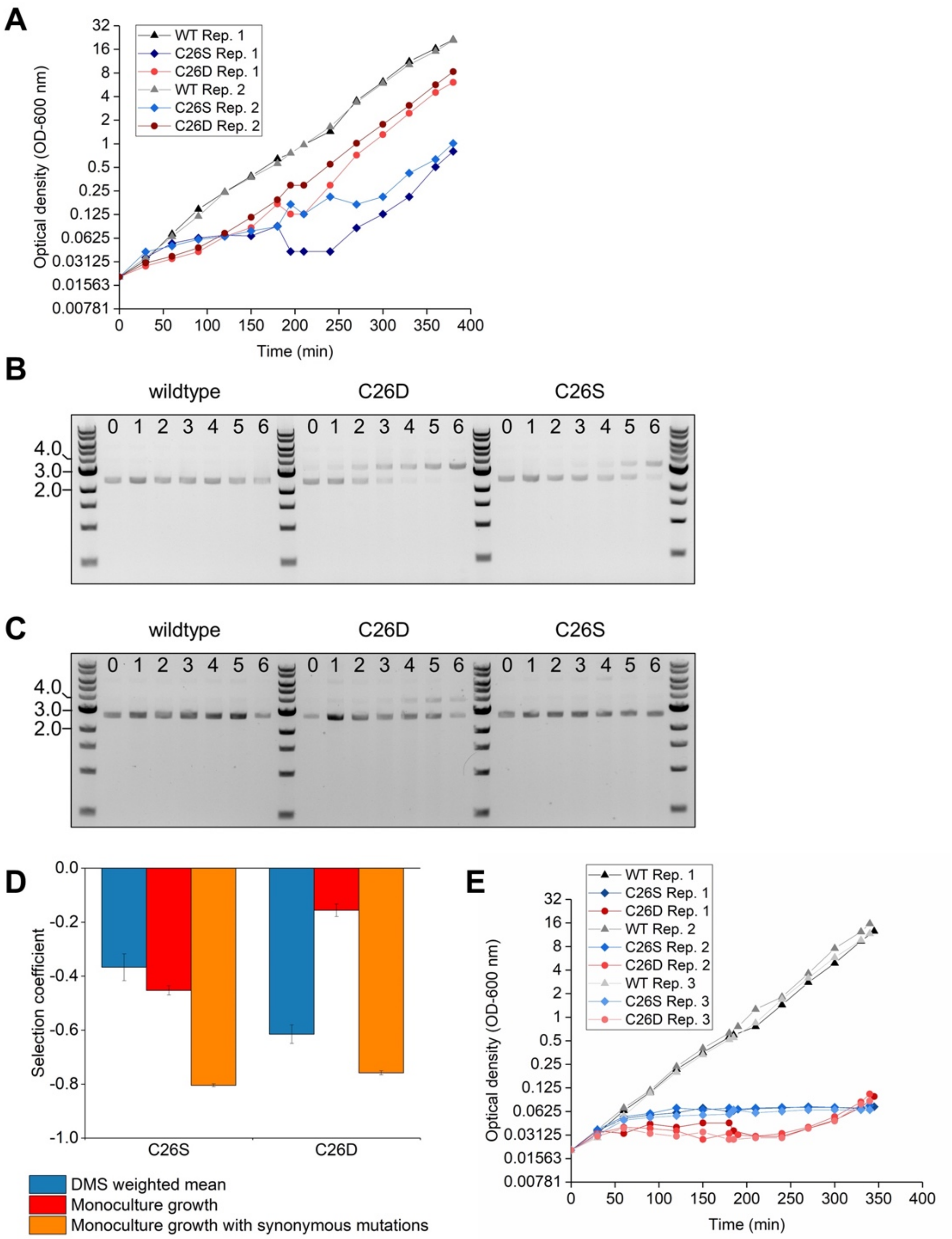


**Figure S10. Correction for transposase insertion in NDM-1.** (**A**) Growth curves of cells expressing unmutated, C26S, and C26D NDM-1 showing how the two mutants' growth accelerated after an initial slower growth period. Sequencing of the *aadB* gene at the end (but not the beginning) of the growth period indicated insertion of an IS-4-like ISVsa5 gene in *aadB*. We collected plasmid extract from every hour of a six-hour induced growth phase to determine the timescale for the transposase insertion in (**B**) cells expressing the C26D and C26S mutants of *NDM-1* and in (**C**) cells expressing the same mutants but with synonymous mutations at L23 and S24 to disrupt the transposase insertion target site. Gels show supercoiled DNA. The lower band represents the pSKunk-1-NDM-1 plasmid while the upper band develops after the transposase insertion occurs. Numbering along the top indicates the hour in the growth experiment from which the DNA was collected. (**D**) Comparison of the selection coefficient for cells with the C26S and C26D mutations as measured by DMS (blue) and monoculture growth (red and orange). The orange represents cells that also have the synonymous mutations at L23 and S24. (**E**) Growth curves of cells expressing unmutated, C26S, and C26D NDM-1 when the gene also contains the synonymous mutations at L23 and S24.

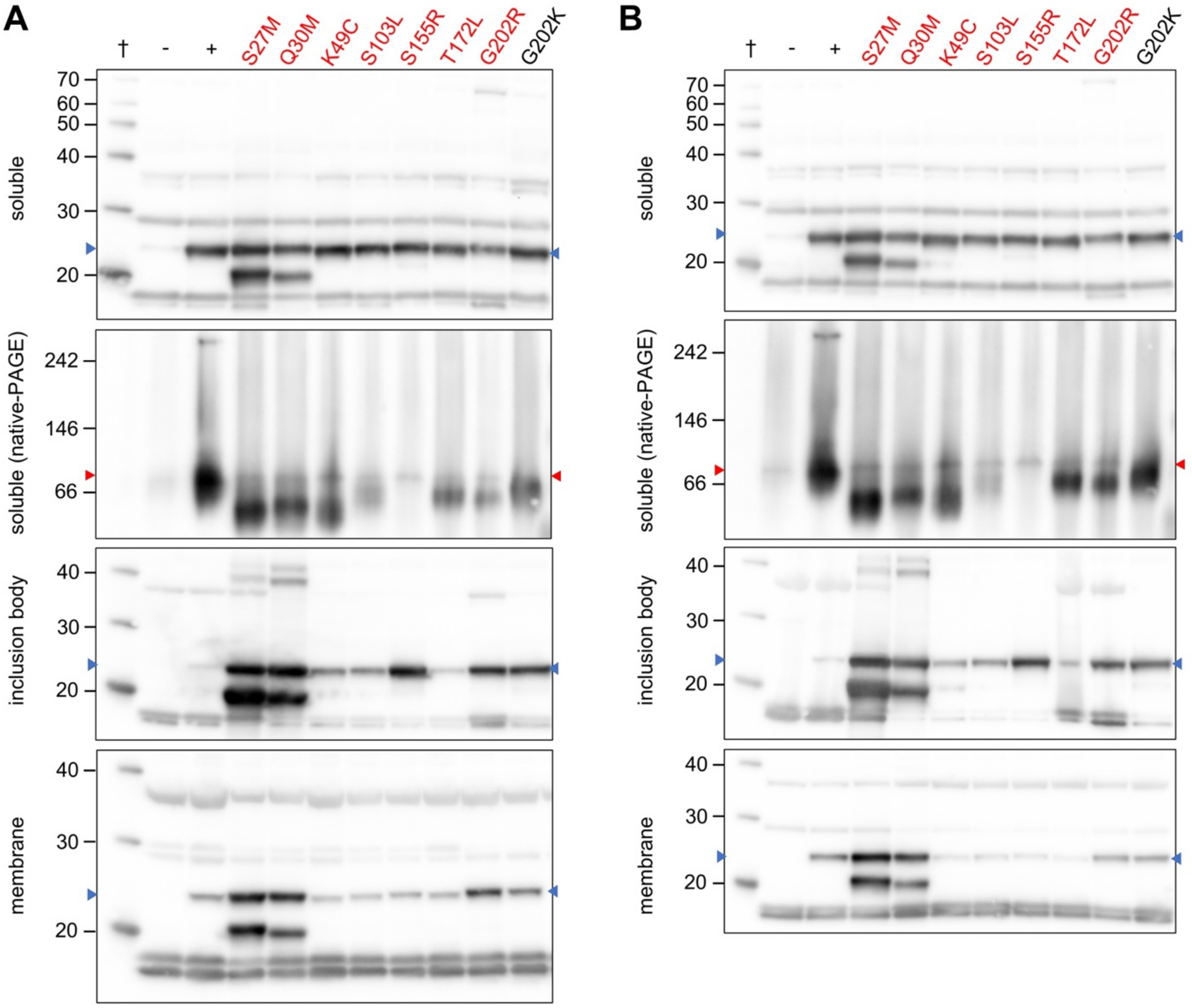


**Figure S11. Western blots of fractionated cells expressing mutants of CAT-I.** Cells expressing CAT-I over six hours of induced growth were harvested, lysed, and fractionated into total soluble, inclusion body, and membrane fractions. Western blots for the (**A**) first and (**B**) second biological replicate are shown with deleterious mutations labeled in red and neutral mutations labeled in black. Samples from fractionated cells without induction of wild-type CAT-I (-) and cells expressing wild-type CAT-I (+) are included on the left of each blot along with the molecular weight standard (†). The expected sizes of the monomer (blue arrowhead) and trimer (red arrowhead) are shown.

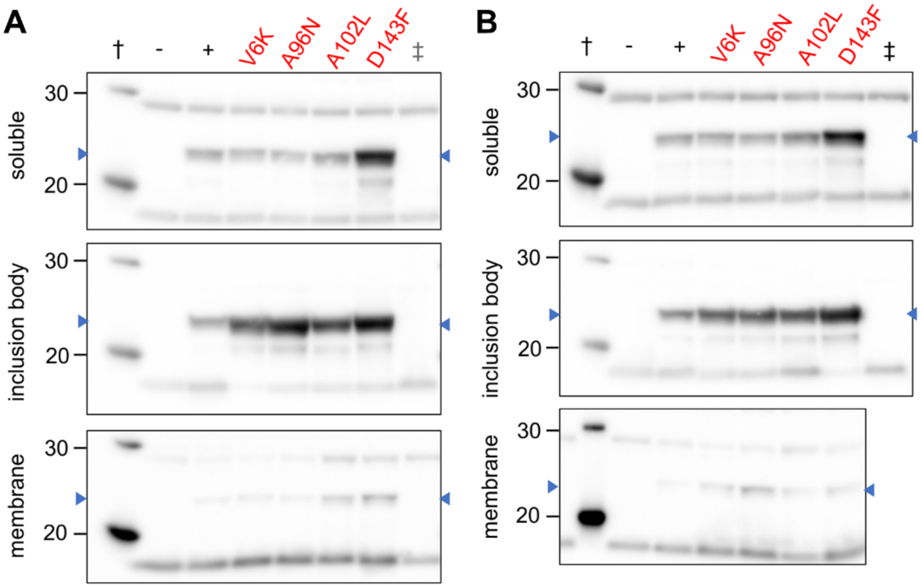


**Figure S12. Western blots of fractionated cells expressing mutants of AadB.** Cells expressing AadB over six hours of induced growth were harvested, lysed, and fractionated into total soluble, inclusion body, and membrane fractions. Western blots for the (**A**) first and (**B**) second biological replicate are shown with deleterious mutations labeled in red and neutral mutations labeled in black. Samples from fractionated cells without induction of wild-type AadB (-) and cells expressing wild-type AadB (+) are included on the left of each blot along with the molecular weight standard (†). CAT-I samples (‡) corresponding to the respective fraction were included on Western blots to identify bands for *E. coli* proteins cross-reacting with the anti-AadB antibodies.

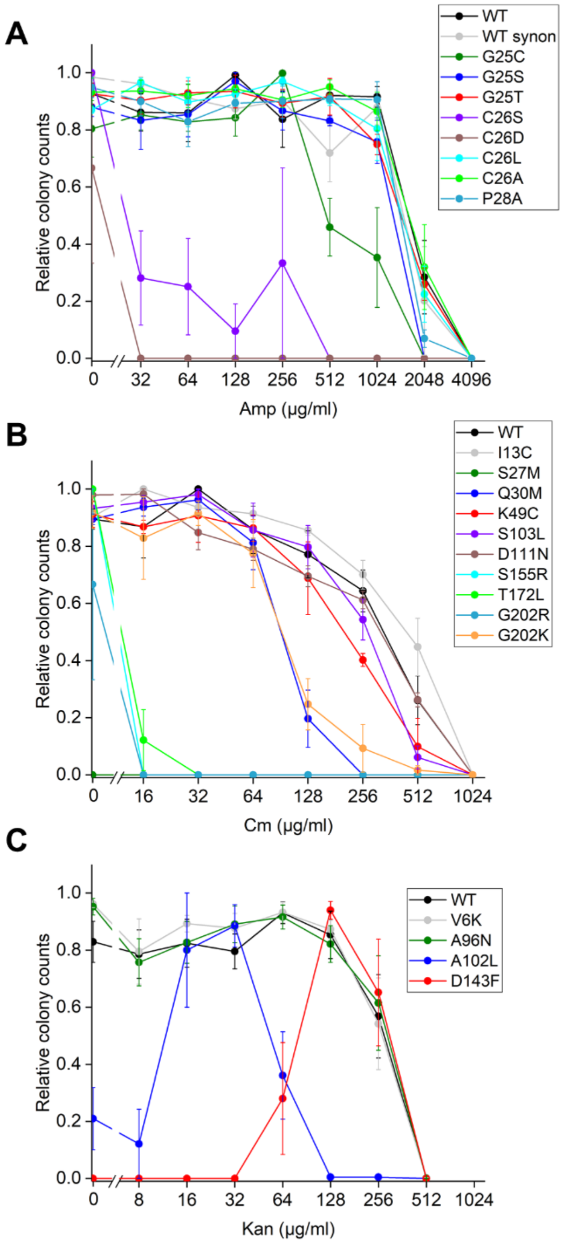


**Figure S13. Relative colony counts for MIC assay.** Relative colony counts for cells expressing wild-type or mutants of (**A**) *NDM-1* plated on ampicillin-containing plates, (**B**) *CAT-I* plated on chloramphenicol-containing plates, and (**C**) *aadB* plated on kanamycin-containing plates. Relative colony counts were calculated from the number of colony forming units (CFU) on a plate divided by the maximum number of CFU at any antibiotic concentration for that same variant. The resulting values were plotted as the mean with error bars representing the standard error of the mean (n = 3 for *NDM-1* and *CAT-I*; n = 5 for *aadB*). Expression of C26S and C26D of NDM-1 along with S27M of CAT-I were lethal in one or more replicates of the MIC assay as no colonies formed on plates lacking the corresponding antibiotic after overnight incubation with the IPTG inducer. Tabulated colony counts are provided as **supplementary data S5**, Supplementary Material online.

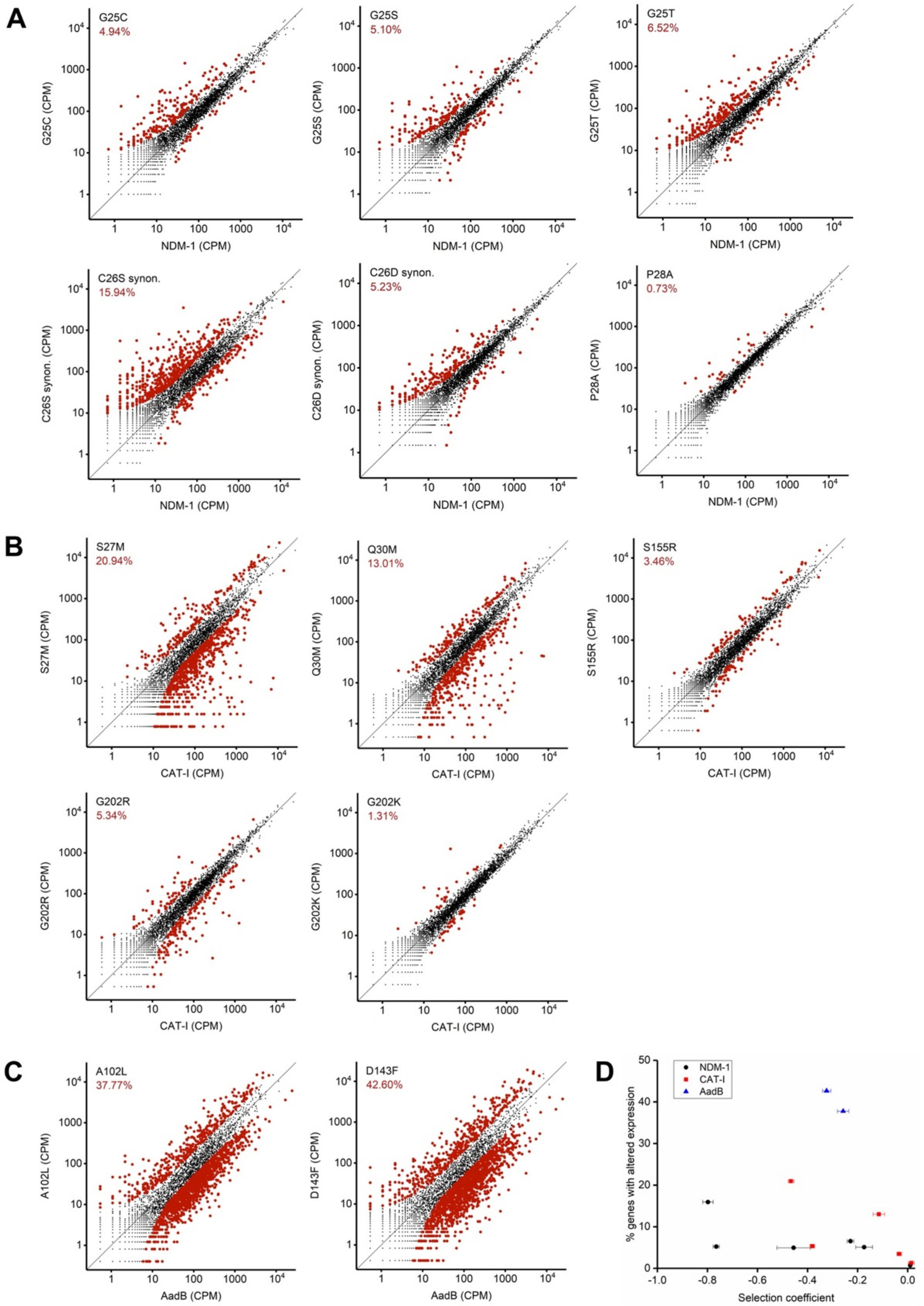


**Figure S14. Differential gene expression caused by missense mutations.** RNA-Seq results showing the gene expression levels in *E. coli* cells expressing a mutant relative to cells expressing the wild-type protein across mutations in (**A**) *NDM-1* (**B**) *CAT-I*, and (**C**) *aadB*. Data indicated with a larger, red circle represents genes in which expression changed by more than twofold with *P*<0.001 significance (*Z*-test) relative to wild-type. The percentage of genes which meet these criteria is indicated as a percentage in red on each plot. CPM, counts per million. (**D**) Relationship between selection coefficient and percentage of genes with more than twofold change in expression and *P*<0.001 significance.

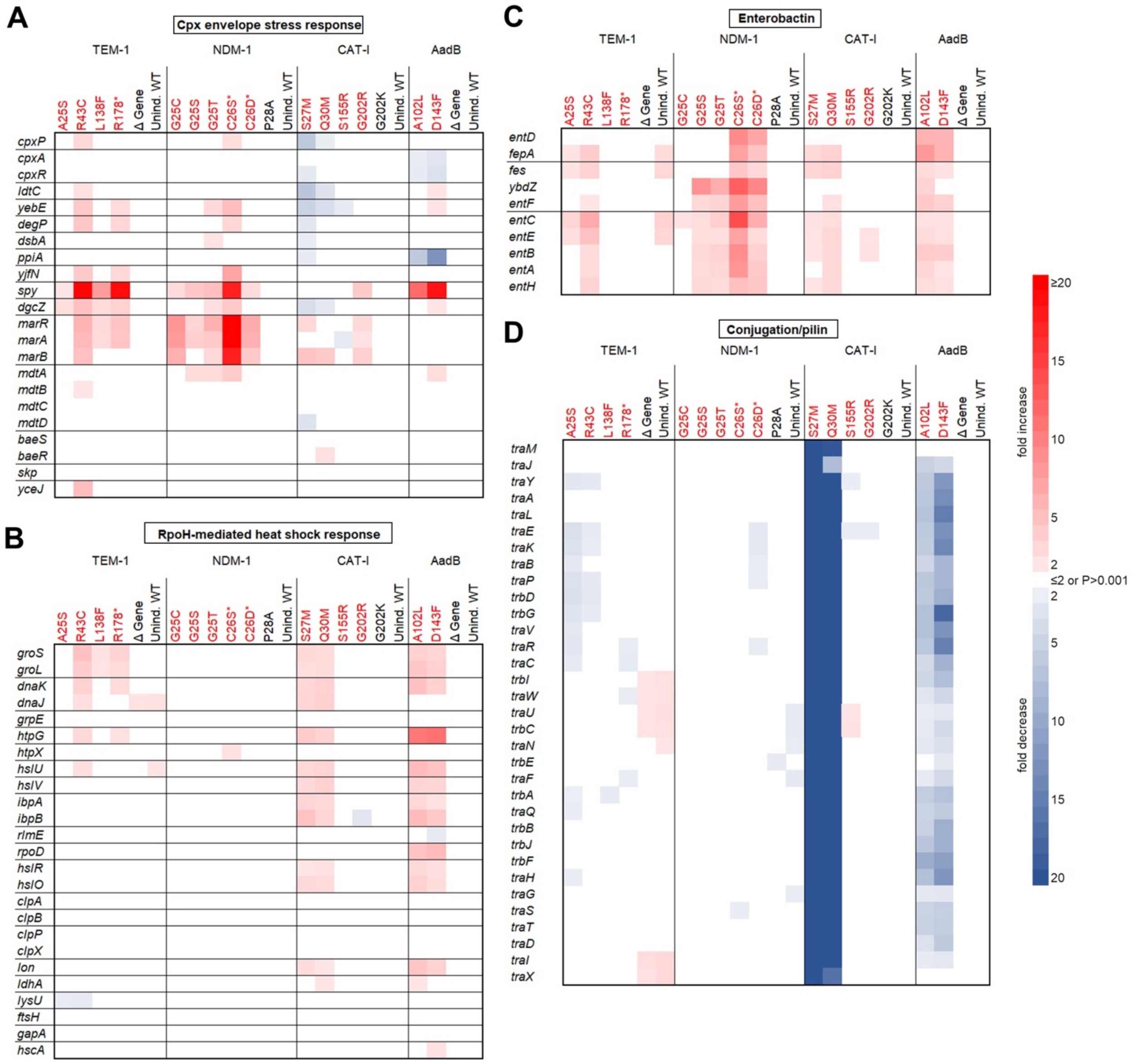


**Figure S15. Effect of mutations related to stress responses and episome.** RNA-Seq experiments revealed differential gene expression for genes corresponding to (**A**) the Cpx envelope stress response, (**B**) the σ$^H$-mediated heat stress response, (**C**) enterobactin biosynthesis, (**D**) and conjugation. Heat maps show genes that differed by greater than twofold expression and *P*<0.001 (Z-test) relative to cells expressing the wild-type antibiotic resistance genes. The color of the mutation indicates whether it is deleterious (red) or neutral (black). Control samples include ones in which wildtype gene expression was not induced (Unind. WT) and induced ones which lack the gene (Δ Gene).

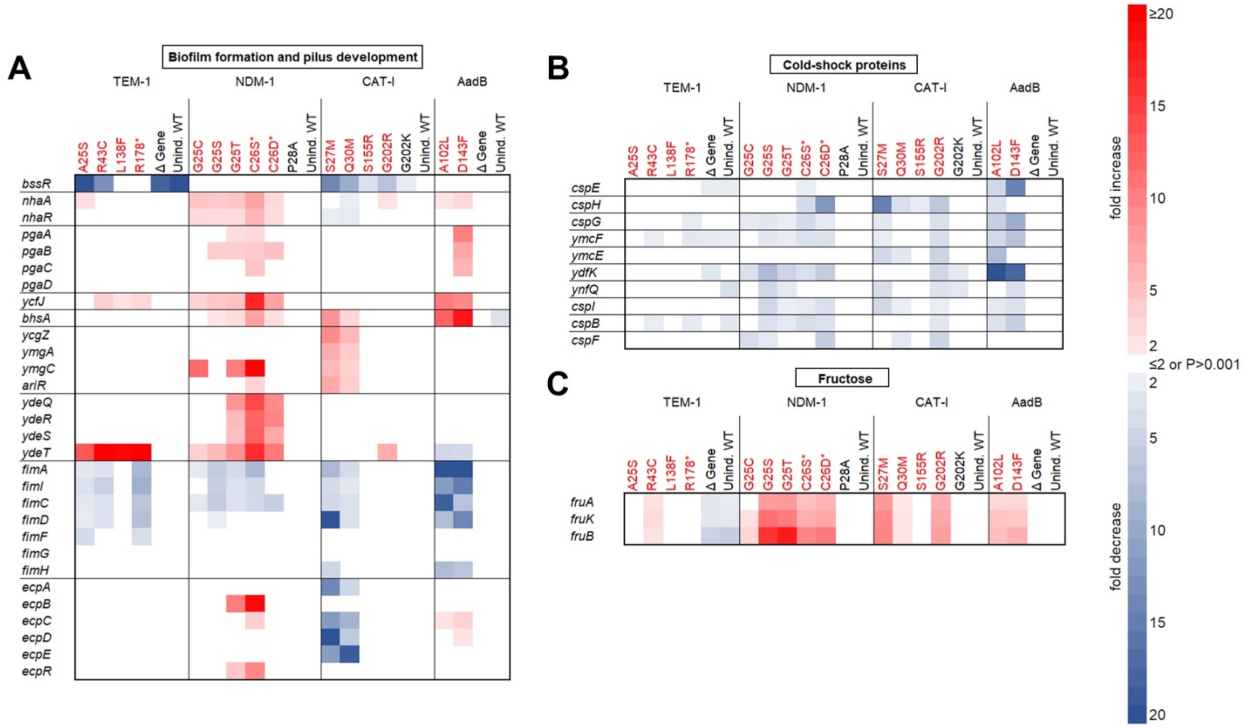


**Figure S16. Effect of mutations on noted operons and pathways.** We identified additional operons and pathways in which multiple genes were differentially expressed relative to their level in cells expressing the wild-type form of the antibiotic resistance gene. Heat maps for select genes belonging to (**A**) biofilm formation and pilus development, (**B**) the cold-shock response, and (**C**) the fructose catabolic process that differed by greater than twofold expression and *P*<0.001 (Z-test) relative to cells expressing the wild-type antibiotic resistance genes. The color of the mutation indicates whether it is deleterious (red) or neutral (black). Control samples include ones in which wildtype gene expression was not induced (Unind. WT) and induced ones which lack the gene (∆ Gene).

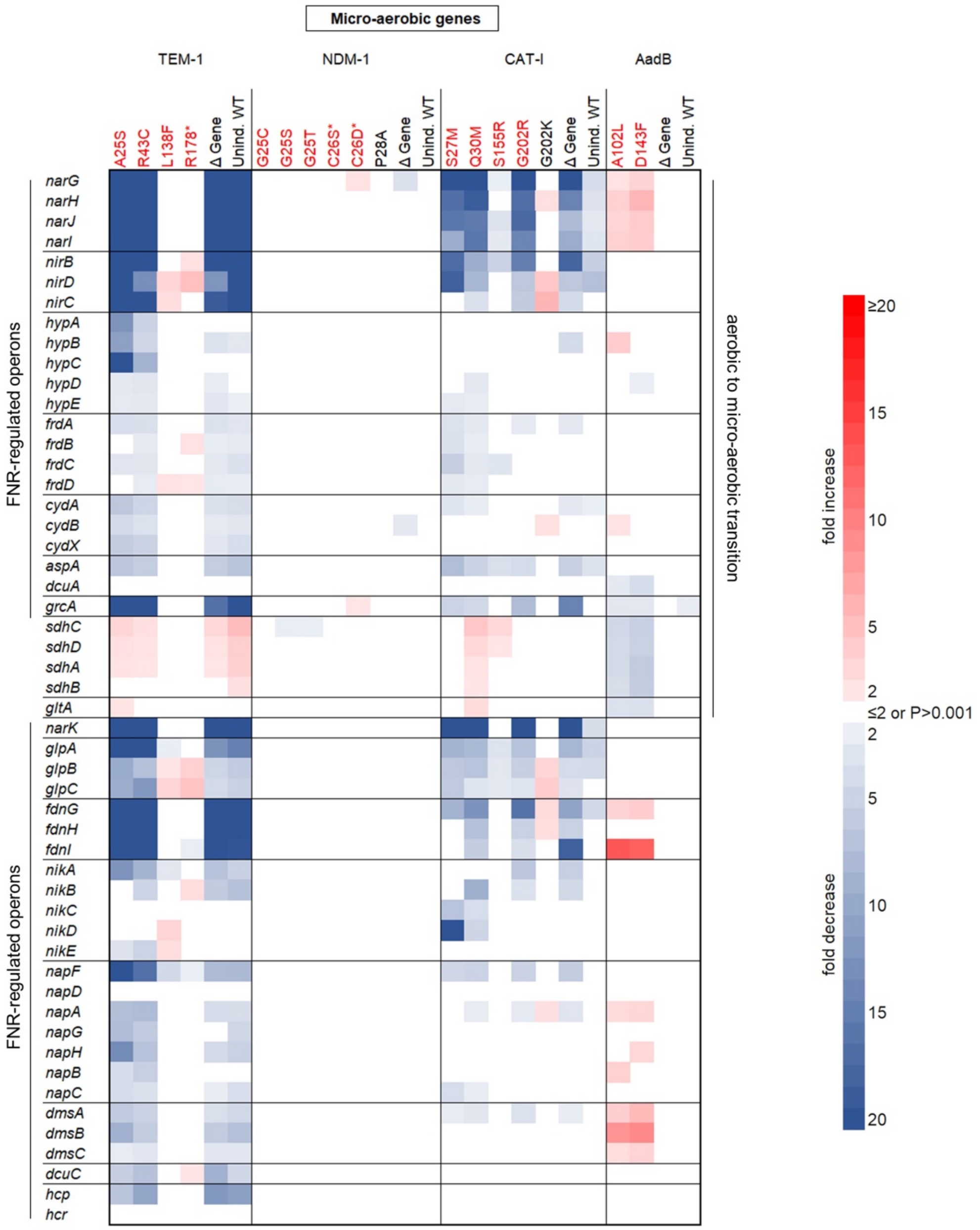


**Figure S17. Differential expression of select genes involved in anaerobic growth.** From RNA-Seq experiments, we also identified a set of genes corresponding to the transition from aerobic to micro-aerobic or anaerobic growth. Heat maps show fold-difference of twofold or more (*P*<0.001, Z-test) in gene expression relative to cells expressing the wild-type form of the antibiotic resistance protein. Genes belonging to operons regulated by FNR are identified along the left (Constantinidou et al. 2006). Genes associated with the transition from aerobic to micro-aerobic growth conditions are labeled on the right (Partridge et al. 2007). The color of the mutation indicates whether it is deleterious (red) or neutral (black). Control samples include ones in which wildtype gene expression was not induced (Unind. WT) and induced ones which lack the gene (Δ Gene).

**Table S1.** Translation initiation rates from methionine substitution in *CAT-I* as predicted by an algorithm described in Reis *et al*. (Reis and Salis 2020). The corresponding weighted mean collateral fitness effect measured by our two replicates of DMS is shown alongside with a white-red gradient to indicate the magnitude of the deleterious fitness effects.

| | Translation Initiation Rate (au) | Fitness |
|---|---|---|
| WT start codon (M1) | 558 | 1.000 |
| F22M | 33 | 0.997 |
| E23M | 704 | 0.975 |
| A24M | 85 | 0.995 |
| F25M | 63 | 1.005 |
| Q26M | 1467 | 0.697 |
| S27M | 811 | 0.610 |
| V28M | 124 | 0.986 |
| A29M | 1212 | 0.697 |
| Q30M | 432 | 0.799 |
| C31M | 30 | 0.997 |
| T32M | 38 | 0.995 |
| Y33M | 29 | 1.004 |
| N34M | 19 | 0.995 |

**Table S2.** Minimum inhibitory concentration (MIC) values in each replicate of the MIC assay[a].

| | Sample | MIC Replicate 1 (µg/ml) | MIC Replicate 2 (µg/ml) | MIC Replicate 3 (µg/ml) | MIC Replicate 4 (µg/ml) | MIC Replicate 5 (µg/ml) |
|---|---|---|---|---|---|---|
| NDM-1 | WT | 4096 | 4096 | 4096 | - | - |
| | WT (synon.) | 2048 | 4096 | 2048 | - | - |
| | G25C | 2048 | 2048 | 1024 | - | - |
| | G25S | 2048 | 2048 | 2048 | - | - |
| | G25T | 4096 | 4096 | 4096 | - | - |
| | C26S | 0 | 0 | 0 | - | - |
| | C26D | 0 | 0 | 0 | - | - |
| | C26L | 4096 | 2048 | 4096 | - | - |
| | C26A | 4096 | 4096 | 4096 | - | - |
| | P28A | 2048 | 2048 | 4096 | - | - |
| CAT-I | WT | 1024 | 1024 | 1024 | - | - |
| | I13C | 1024 | 1024 | 1024 | - | - |
| | S27M | 0 | 0 | 0 | - | - |
| | Q30M | 128 | 256 | 256 | - | - |
| | K49C | 512 | 1024 | 512 | - | - |
| | S103L | 512 | 1024 | 512 | - | - |
| | D111N | 1024 | 1024 | 1024 | - | - |
| | S155R | ≤16 | ≤16 | ≤16 | - | - |
| | T172L | ≤16 | ≤16 | 32 | - | - |
| | G202R | ≤16 | ≤16 | 0 | - | - |
| | G202K | 256 | 512 | 256 | - | - |
| AadB | WT | 512 | 512 | 512 | 256 | 512 |
| | V6K | 512 | 512 | 512 | 256 | 512 |
| | A96N | 512 | 512 | 512 | 256 | 512 |
| | A102L | 128 | 64 | 128 | 64 | 128 |
| | D143F | 512 | 512 | 512 | 256 | 512 |

[a] The antibiotics used were ampicillin (*NDM*-1), chloramphenicol (*CAT*-I), and kanamycin (*aadB*). The C26S, C26D, and WT (synon.) samples also contain the synonymous mutations at L23 and S24 that disrupt the transposase insertion target site. The lowest chloramphenicol concentration tested was 16 µg/ml.

**Supplementary References**

Constantinidou C, Hobman JL, Griffiths L, Patel MD, Penn CW, Cole JA, Overton TW. 2006. A reassessment of the FNR regulon and transcriptomic analysis of the effects of nitrate, nitrite, NarXL, and NarQP as Escherichia coli K12 adapts from aerobic to anaerobic growth. *J. Biol. Chem.* 281:4802–4815.

Kram KE, Henderson AL, Finkel SE. 2020. Escherichia coli has a unique transcriptional program in long-term stationary phase allowing identification of genes important for survival. *mSystems* 5.

Partridge JD, Roberts RE, Poole RK. 2007. Transition of Escherichia coli from Aerobic to Micro-aerobic Regulatory Components involves fast and slow reacting regulatory components. *J. Biol. Chem.* 282:11230-11237.

Reis AC, Salis HM. 2020. An automated model test system for systematic development and improvement of gene expression models. *ACS Synth. Biol.* 9:3145–3156.

**Captions for data files**

**Data S1. Excel file for DMS sequencing counts, fitness values, and associated statistics for collateral fitness effects in NDM-1.**

**Data S2. Excel file for DMS sequencing counts, fitness values, and associated statistics for collateral fitness effects in CAT-I.**

**Data S3. Excel file for DMS sequencing counts, fitness values, and associated statistics for collateral fitness effects in AadB.**

**Data S4. Excel file for RNA-seq sequencing counts, fold-difference in expression, and associated statistics.**

**Data S5. Excel file for NDM-1, CAT-I, and AadB MIC assay colony counts.**
